# eIF3 mRNA selectivity profiling reveals eIF3k as a cancer-relevant regulator of ribosome content

**DOI:** 10.1101/2022.08.28.505560

**Authors:** Haoran Duan, Siqiong Zhang, Yoram Zarai, Rupert Öllinger, Yanmeng Wu, Li Sun, Cheng Hu, Guiyou Tian, Roland Rad, Yabin Cheng, Tamir Tuller, Dieter A. Wolf

**Affiliations:** State Key Laboratory of Stress Biology and Fujian Provincial Key Laboratory of Innovative Drug Target Research, School of Pharmaceutical Sciences, Xiamen University, Xiamen, China; Department of Biomedical Engineering, Tel Aviv University, Tel Aviv 6997801, Israel; Institute of Molecular Oncology and Functional Genomics and Center for Translational Cancer Research (TranslaTUM), School of Medicine, Technical University of Munich, Munich 81675, Germany; Ganzhou Key Laboratory for Drug Screening and Discovery, School of Geography and Environmental Engineering, Gannan Normal University, Ganzhou 341000 Jiangxi, China; German Cancer Consortium (DKTK), German Cancer Research Center (DKFZ), Heidelberg, Germany; Department of Internal Medicine II, Klinikum rechts der Isar, Technical University Munich, Munich 81675, Germany; The Sagol School of Neuroscience, Tel-Aviv University, Tel Aviv 6997801, Israel

## Abstract

eIF3, whose subunits are frequently overexpressed in cancer, regulates mRNA translation from initiation to termination, but mRNA-selective functions of individual subunits remain poorly defined. Using multi-omic profiling upon acute depletion of eIF3 subunits, we observed that while eIF3a, b, e, and f markedly differed in their impact on eIF3 holo-complex formation and translation, they were each required for cancer cell proliferation and tumor growth. Remarkably, eIF3k showed the opposite pattern with depletion promoting global translation, cell proliferation, tumor growth, and stress resistance through repressing the synthesis of ribosomal proteins, especially RPS15A. Whereas ectopic expression of RPS15A mimicked the anabolic effects of eIF3k depletion, disruption of eIF3 binding to the 5’-UTR of *RSP15A* mRNA negated them. Supported by mathematical modeling, our data uncovers eIF3k-l as a mRNA-specific module which, through controlling *RPS15A* translation, serves as a rheostat of ribosome content to secure spare translational capacity that can be mobilized during stress.

## INTRODUCTION

Protein synthesis is a complex process requiring tight coordination of discrete steps including the initiation of mRNA translation, translation elongation and termination as well as co-translational folding, subcellular targeting, and quality control of the nascent polypeptides. In canonical initiation, mRNA is recruited to the ribosome through eIF4E-mediated binding of the m7G cap. eIF4E together with eIF4G then recruits to the mRNA a complex consisting of eIF3 and the 40S ribosome loaded with methionyl-tRNA and additional eIFs (Sonenberg and Hinnebusch, 2009). Upon scanning for the start codon, the 60S ribosomal subunit joins to produce an actively translating 80S ribosome.

Apart from canonical translation initiation via the 5’-cap, the 13-subunit eIF3 complex mediates several non-canonical modes of initiation through direct interaction with mRNAs or mRNA binding proteins (Wolf et al., 2020). Using PAR-CLIP, eIF3 was shown to interact with structured 5’-UTR motifs of hundreds of mRNAs, resulting in translational activation or repression (Lee et al., 2015). For example, phosphorylation-controlled binding of eIF3 to c-Jun mRNA was shown to regulate its cap-dependent but eIF4E-independent function in translation (Lamper et al., 2020; Lee et al., 2016). Likewise, transient recruitment of eIF3 to their 3’-UTRs promotes a burst in the translation of mRNAs encoding T cell receptors (De Silva et al., 2021). Conversely, binding of eIF3 to the ferritin light chain mRNA led to inhibition of its translation (Pulos-Holmes et al., 2019). In addition, eIF3 binding sites in mRNAs frequently overlap with sites of N6 methyl adenosine (m6A) modification, and eIF3 was shown to bind to m6A either directly or through m6A reader and writer proteins (Meyer et al., 2015; Wolf et al., 2020). Thus, eIF3 is positively as well as negatively involved in many different forms of canonical and non-canonical translation initiation.

However, not all eIF3 subunits are essential for protein synthesis, cell growth, and survival (Wagner et al., 2016), suggesting that some have mRNA selective functions that underlie eIF3’s multifunctionality. For example, we have reported that subunits eIF3d and eIF3e constitute a module within the eIF3 holo-complex that selectively promotes the synthesis of proteins with membrane-associated functions, including subunits of the mitochondrial electron transport chain (ETC) (Lin et al., 2020; Shah et al., 2016). Likewise, eIF3f was shown to advance sperm-specific translation through interaction with proteins that bind MIWI/piRNAs in 3’-UTRs thus leading to circularization of mRNA which is thought to promote translation (Dai et al., 2019). Even more recently, eIF3g was implicated in neuron-specific translation in C. elegans (Blazie et al., 2021). Furthermore, eIF3c was shown to be dispensable for global translation and viability in mice but to direct the translation of the Sonic Hedgehog receptor PTCH1 through a pyrimidine-rich motif in the 5’-UTR (Fujii et al., 2021). Finally, the eIF3k/l module was reported to be dispensable for viability in C. elegans, whereas its removal enhances stress resistance, although this effect remains to be attributed to mRNA-selective changes in translation (Cattie et al., 2016). To date, no study has assessed the impact on holo-complex formation and mRNA-specific translation of individually disabling multiple eIF3 subunits in the same system.

In terms of human physiology, many eIF3 subunits are overexpressed in cancer and can drive de novo holo-complex formation and increased protein synthesis along with cell transformation in vitro (Hershey, 2015). However, despite striking progress in the structural analysis of the eIF3 complex (Eliseev et al., 2018; des Georges et al., 2015; Hashem et al., 2013; Simonetti et al., 2016; Thoms et al., 2020), mechanistic insight into the mRNA selective functions of eIF3 subunits remains scarce and no drugs exist to target eIF3 in cancer or other diseases.

Testing the concept that eIF3 consists of distinct functional modules, we have embarked on a systematic loss-of-function study to evaluate mRNA-selectivity of module-defining eIF3 subunits in cell proliferation and cancer growth. While eIF3a, b, e, and f are found to promote cell proliferation and tumor growth despite regulating distinct sets of mRNAs, eIF3k limits cell and tumor growth by repressing global translation. Mechanistically, this eIF3k-dependent repression is mediated by binding of eIF3k to the 5’-UTR of the mRNA encoding ribosomal protein RPS15A. Our studies define eIF3 as a multi-functional complex consisting of distinct modules imposing positive as well as negative regulation on mRNA-selective translation with the eIF3k-l module serving as a rheostat of ribosome content.

## RESULTS

### Differential requirement of eIF3 subunits for eIF3 holo-complex formation

To assess subunit-specific functions of eIF3, we generated HCT116 colon cancer cell lines in which individual eIF3 subunits can be rapidly and reversibly depleted by small molecule-induced degradation. We targeted the following eIF3 subunits (**Figure 1A**): (i.) eIF3a, a core subunit known to nucleate the assembly of the holo-complex (Smith et al., 2016; Wagner et al., 2016), (ii.) eIF3b, a subunit of the functionally distinct “b-g-i” module, (iii.) eIF3f, a subunit which was reported to inhibit rather than activate translation (Aguero et al., 2017; Shi et al., 2006), (iv.) eIF3e, a subunit of the “d-e” module involved in the synthesis of membrane-associated proteins (Lin et al., 2020; Shah et al., 2016), and (v.) eIF3k, a subunit of the non-essential “k-l” module, which has cryptic mRNA cap-binding activity through eIF3l (Kumar et al., 2016). We used CRISPR-mediated genome editing to add the auxin-inducible degradation domain (mAID) to both alleles of eIF3a, b, e, f, and k (Natsume et al., 2016). In addition, both alleles of each gene were modified with either mClover or mCherry for allele-specific protein imaging (**Figure EV1A**). Fluorescence microscopy revealed predominantly cytoplasmic localization of all five eIF3 subunits (**Figure EV1B**), a pattern which is consistent with their established role in cytoplasmic mRNA translation. Thus, epitope tagging with mAID and fluorescent proteins does not seem to interfere with normal subcellular localization of eIF3 subunits.

**Figure 1.**
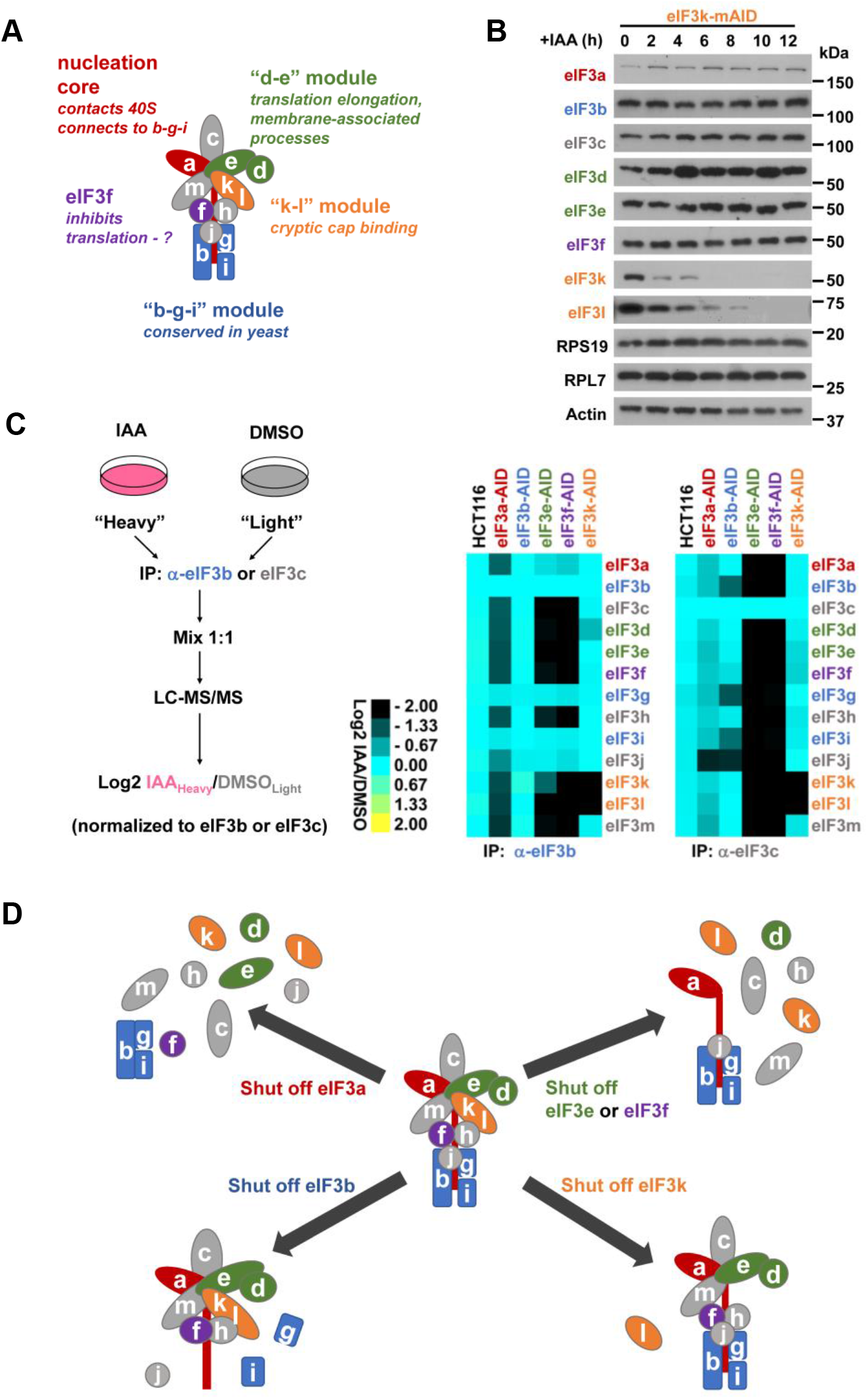
Effect of conditional depletion of eIF3 subunits on the eIF3 holo-complex. **(A)** Schematic of holo-eIF3 with the five subunits targeted for conditional depletion highlighted in different colors. **(B)** HCT116 cells in which eIF3k was homozygously modified with the mAID domain to trigger auxin-dependent degradation were exposed to 500 μM indole-3-acetic acid (IAA) for the periods shown, and cell lysate was subjected to immunoblotting with the indicated antibodies. **(C)** Analysis of holo-eIF3 complex formation by quantitative SILAC proteomics. eIF3-mAID cell lines were grown either in regular “light” (L) media or in media supplemented with stable “heavy” (H) lysine and arginine. Heavy-labelled cells were treated with IAA for 12 hours to shut off the expression of eIF3 subunits. eIF3 was immunopurified from both conditions using antibodies against eIF3b or eIF3c. Upon trypsin digestion, corresponding L (DMSO vehicle control) and H (IAA) samples were mixed and analyzed by LC-MS/MS (left panel). Average changes in H/L ratios were determined, normalized to the bait (eIF3b or eIF3c, respectively), and plotted in a heatmap (right panel). **(D)** Graphical summary of the effect of eIF3 subunit depletion on holo-eIF3 as determined by quantitative proteomics and native PAGE analysis (Figure S2B).

Unlike RNAi-mediated depletion of eIF3 subunits for prolonged duration (e.g. 3 days (Wagner et al., 2016)) and potentially marked by variable efficiency and off-target activity, auxin-induced degradation led to acute depletion thus minimizing adaptive effects expected after long-term depletion. For example, eIF3b-mAID and eIF3k-mAID were selectively downregulated within 2 hours of auxin (IAA) addition whereas the expression of the remaining 7 eIF3 subunits tested was not majorly affected (**Figure 1B, EV1C**). eIF3a-mAID and eIF3f-mAID showed a similar depletion but also led to co-depletion of eIF3k (**Figure EV1C**). Depletion of eIF3e-mAID caused co-depletion of eIF3d and eIF3k as previously observed (Shah et al., 2016; Wagner et al., 2016; Yen and Chang, 2000). Addition of IAA to parental HCT116 cells had no effect on the expression of eIF3 subunits.

We next determined the consequences of depleting eIF3 subunits for eIF3 holo-complex formation by quantitative SILAC proteomics (Cox et al., 2009). For this, we maintained eIF3-mAID cell lines either in regular “light” (L) media or in media supplemented with stable “heavy” (H) lysine and arginine. Heavy-labelled cells were treated with IAA for 12 hours to shut off the expression of eIF3 subunits. eIF3 was immunopurified from both conditions using antibodies against eIF3b or eIF3c. Upon trypsin digestion, corresponding L (DMSO control) and H (IAA) samples were mixed and analyzed by LC-MS/MS (**Figure 1C**). The data showed that acute depletion of eIF3a led to a major disassembly of holo-eIF3 while largely sparing the eIF3b-g-i sub-complex (**Figure 1C, D, EV2A**). Conversely, depletion of eIF3b led to a substantial depletion of the eIF3b-g-i sub-complex, while mostly maintaining the octameric core complex. Depletion of eIF3e affected holo-eIF3 formation identically to depletion of eIF3f, leaving intact a residual complex consisting of eIF3a, b, g, i, and j, which closely resembles the essential core complex present in S. cerevisiae (Valášek et al., 2017). Lastly, depletion of eIF3k maintained an eIF3 complex that only lacked the eIF3k-l module (**Figure 1C,D, EV2A**).

The conclusions drawn from the quantitative proteomic analysis were qualitatively confirmed by native PAGE analysis (**Figure EV2B**). Here again, depletion of eIF3a led to a stark reduction in the high molecular weight complex co-migrating with holo-eIF3 in parental HCT116 cells as measured with antibodies against eIF3b, c, e, f, k, i, and g. However, in line with the SILAC data, a smaller complex representing the “b-g-i” sub-complex was maintained (**Figure EV2B**). In contrast, depletion of eIF3k did not appreciably affect the apparent eIF3 holo-complex, a finding that is consistent with the minor impact of loss of the eIF3k-l module on the total mass of the complex. Finally, cells depleted of eIF3e and eIF3f retained substantial amounts of the core complex containing eIF3a, b, g, and i (**Figure EV2B**).

In summary, the data obtained with acute depletion of eIF3 subunits showed that removal of individual eIF3 subunits distinctly reshapes eIF3 complex architecture rather than entirely destroying the complex (**Figure 1D**). Thus, eIF3 appears to be organized in distinct subunit-specific functional modules.

### Subunit-specific functions of eIF3 in cell proliferation and tumor growth

To begin to assess module-specific functions, we tested the effect of acute depletion of eIF3 subunits on cell proliferation. Whereas eIF3a, b, e, and f were equally required for cell proliferation as determined by a surrogate metabolic assay measuring NAD(P)H levels (MTT assay) (**Figure 2A**) as well as colony formation (**Figure 2B, EV3A**), eIF3k was entirely dispensable for both. In fact, cells depleted of eIF3k showed higher proliferation than the syngeneic controls maintained in the absence of IAA (**Figure 2A, B**). The rapid depletion system also allowed us to determine the effect of eIF3 subunits on cell cycle dynamics. Re-entry into S phase upon release from a block in G0/G1 induced by serum starvation was greatly impaired in cell lines depleted of eIF3a, b, and e. Whereas depletion of eIF3f had a milder effect on cell cycle re-entry, cells deficient in eIF3k did not show a difference in cell cycle kinetics from cells maintained in the absence of IAA (**Figure 2C**).

**Figure 2.**
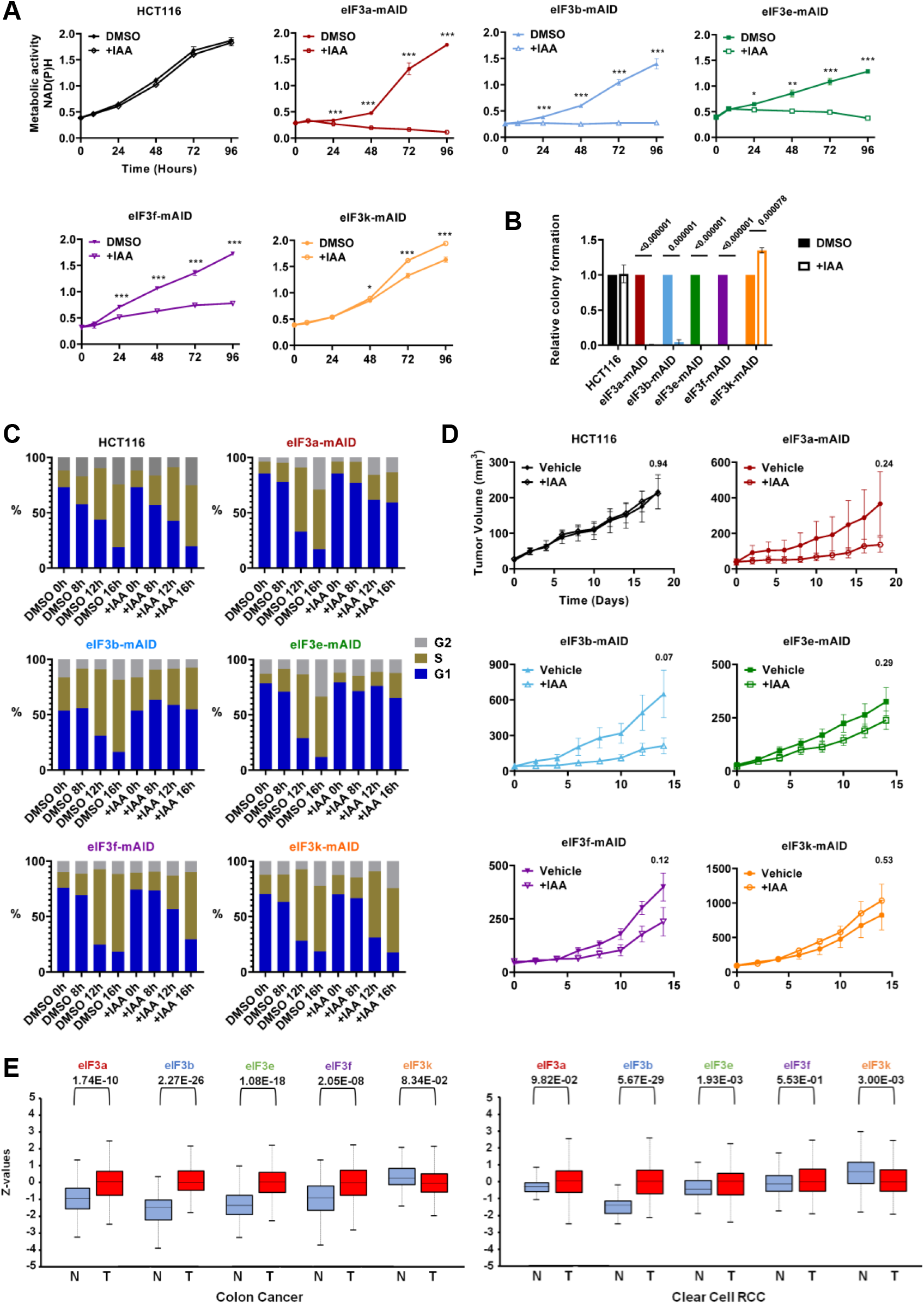
Effect of depletion of eIF3 subunits on cell and tumor growth. **(A)** Parental HCT116 and the five different conditional eIF3-mAID cell lines were treated with the vehicle DMSO or with IAA for the periods indicated. Metabolic activity (i.e. NAD(P)H) as a proxy of cell proliferation was determined by MTT assay. Graphs represent means ± SD, n = 5 - 6. Asterisks denote: * p < 0.005, ** p < 0.00005, *** p < 0.000005 (two-stage step-up method of Benjamini, Krieger, and Yekutieli). **(B)** 1 × 10^3^ cells of the indicated cell lines were plated and grown in DMSO or IAA for 2 weeks. Colony numbers were counted and are shown in a bar graph relative to the DMSO-treated control. Note that very low numbers of colonies were obtained for IAA treated eIF3a-mAID, eIF3e-mAID and eIF3f-mAID - hence no bars are visible. Bars represent means ± SD, n = 3. Numbers indicate p values (unpaired Student’s t-test). **(C)** The indicated cell lines were subjected to serum withdrawal (0 % FBS) for 16 h. DMSO or IAA were added and cells were stimulated to re-enter the cell cycle by re-feeding with media containing 10% FBS. Cells were fixed after the indicated time points, and the cellular DNA content was measured by flow cytometry. Data are representative of 2 (HCT116, eIF3a-mAID, eIF3b-mAID) of 3 (eIF3e-mAID, eIF3f-mAID, eIF3k-mAID) experiments. **(D)** 1 × 10^6^ eIF3-mAID cells were injected into nude mice. When tumors reached a diameter of ∼ 4 mm, mice were treated with vehicle or 500 mg/kg IAA, and tumor growth was measured for 2 weeks. Graphs represent means ± SD, n = 5 to 7. Numbers indicate p values (two-stage step-up method of Benjamini, Krieger, and Yekutieli). **(E)** Expression of the indicated eIF3 proteins in 97 colon adenocarcinomas versus 100 normal colonic mucosas and 110 clear cell renal cell carcinomas versus 84 normal kidney samples. Log2 spectral count data from CPTAC (Chen et al., 2019) via UALCAN portal (Chandrashekar et al., 2022).

The same differential requirement of eIF3 subunits was observed upon growing the conditional depletion cell lines as xenograft tumors in mice. Once tumors reached a diameter of ∼4 mm (volume ∼50 mm^3^), eIF3 subunits were depleted by daily oral gavage of IAA. IAA did not affect the growth of tumors established from parental HCT116 cells relative to the vehicle (**Figure 2D, EV3B**). Whereas tumor sizes increased in mice treated with the vehicle control, the growth of eIF3a-mAID and eIF3b-mAID tumors was substantially reduced upon administration of IAA (**Figure 2D, EV3B**). eIF3e-mAID and eIF3f-mAID tumors were also suppressed by IAA, although to a lesser extent (**Figure 2D, EV3B**). In marked contrast, tumors grew to bigger sizes when eIF3k was depleted (**Figure 2D, EV3B**), suggesting that eIF3k may play a tumor suppressive role, whereas the other eIF3 subunits examined have pro-tumorigenic function. Consistent with this notion are proteomic data showing that average protein levels of eIF3a, b, e, and f were upregulated in human colon and renal cancers, whereas eIF3k protein was downregulated (**Figure 2E**).

### mRNA selectivity of eIF3 subunits revealed by ribonomic and proteomic profiling

To determine the effect of acute depletion of eIF3 subunits on global mRNA translation, we performed sucrose density gradient fractionation to separate actively translating polysomes (P) from 40S, 60S, and 80S monosomes (M). Addition of IAA to the various cell lines for 12 h led to ∼72 – 90% depletion of eIF3 subunits (**Figure 3A, EV4A**). Whereas IAA did not affect the P/M ratio of parental HCT116 cells, depletion of eIF3a, e, and f subunits led to substantial decreases in the P/M ratios indicating substantial inhibition of translation with eIF3a depletion showing the most pronounced effect (**Figure 3B, EV4B**). Addition of IAA to eIF3b-mAID cells for 12 hours did not affect the P/M ratio even so eIF3b was 73% depleted (**Figure 3A, B, EV5A**). Only ∼90% depletion of eIF3b after 24 h of IAA treatment led to a decrease in translation (**Figure EV5A**). In contrast, depletion of eIF3k did not decrease translation but rather caused a 6% (±2.7%) increase in the P/M ratio, a finding consistent with a previous study (Herrmannová et al., 2020) and with the increased cell proliferation and tumor growth observed with the eIF3k-mAID cell line (**Figure 2**).

**Figure 3.**
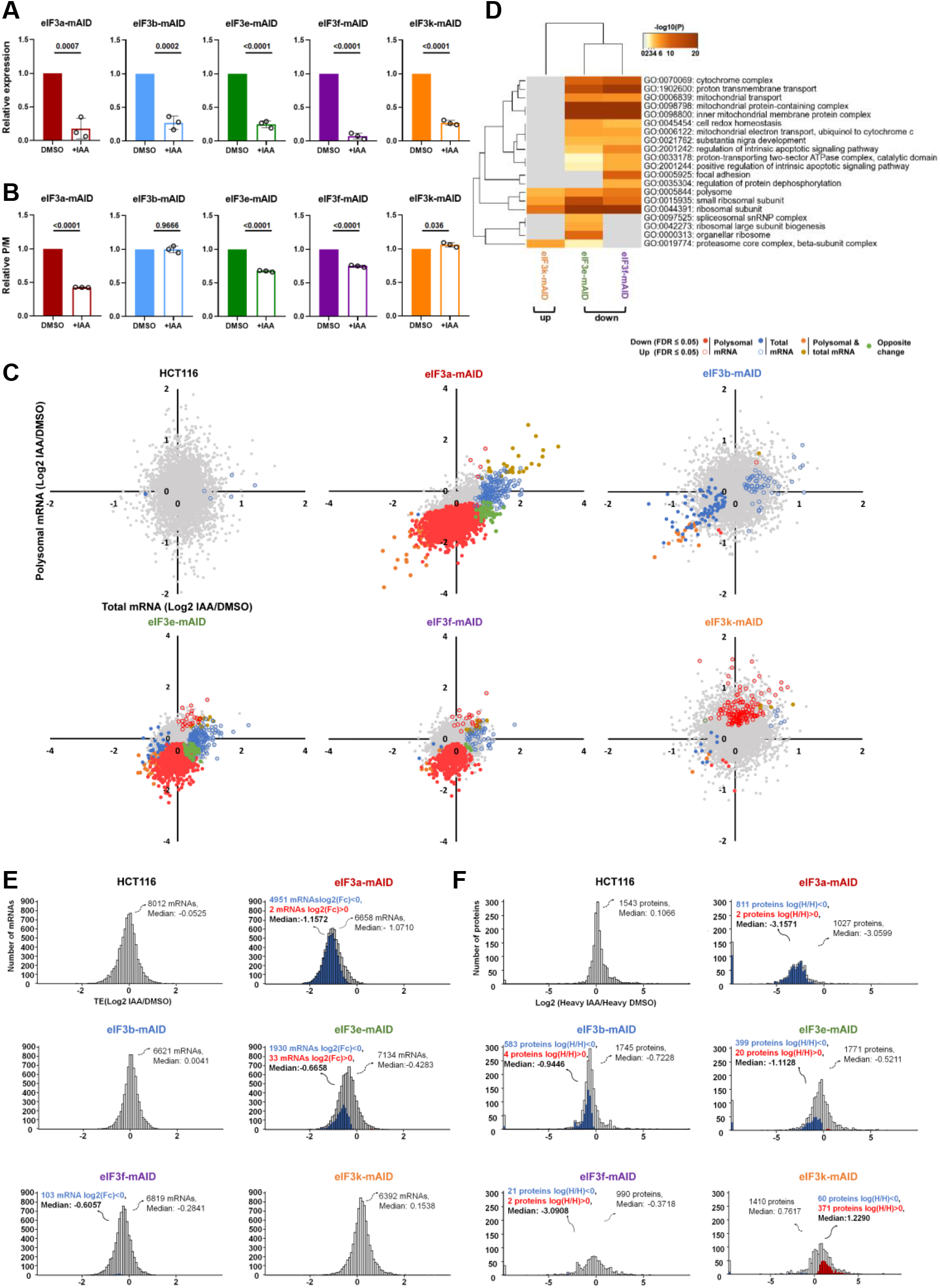
Effect of depletion of eIF3 subunits on mRNA translation. **(A)** The indicated conditional eIF3 cell lines were exposed to 500 μM IAA for 12 h to induce the degradation of the respective eIF3 subunits. Cell lysates were subjected to immunoblotting to quantify the expression of the indicated eIF3 subunits. Signals were normalized to the reference signal of actin and plotted relative to the vehicle DMSO. Bars represent means ± SD, n = 3. Numbers indicate p values (unpaired Student’s t-test). **(B)** The indicated cell lines were exposed to IAA for 12 h, and cell lysates were separated by sucrose density gradient centrifugation. The monosomal (M) and polysomal (P) peaks were quantified, and P/M ratios relative to the vehicle DMSO were plotted. Bars represent means ± SD, n = 3. Numbers indicate p values (unpaired Student’s t-test). **(C)** Scatter plots of changes in polysomal mRNA versus total mRNA upon depletion of the indicated eIF3 subunits. **(D)** Enrichment of Gene Ontology terms in the sets of polysome-associated mRNAs decreased in the polysomal fractions of eIF3f or eIF3e depleted cells or increased in eIF3k depleted cells. **(E)** Distribution histograms of translational efficiencies (TEs) of the mRNA sets shown in (**C**). The total number of mRNAs and the median TE of each dataset are indicated. The number of up and down changes with FDR ≤ 0.2 as well as the median of all changed items are indicated in red, blue, and black bold print. **(F)** Distribution histograms of the protein synthesis ratios determined in eIF3 subunit depleted versus replete cells. The total number of measured proteins and the median synthesis change of each dataset are indicated. The number of up and down changes with FDR ≤ 0.1 as well as the median of all changed items are indicated in red and blue print.

To assess translation of individual mRNAs upon depletion of eIF3 subunits, we performed RNA-seq of triplicate samples of mRNA contained in sucrose density gradient fractions corresponding to > 2 polyribosomes (polysomal mRNA). In parallel, we sequenced triplicate samples of total cellular mRNA and compared the data to the polysomal mRNA-seq (analysis strategy illustrated in **Figure EV4C**, all data in **Supplementary Data Files 1, 2**). As shown in a scatterplot of Log2 fold changes of polysomal mRNA versus Log2 fold changes of total mRNA, IAA treatment for 12 h did not lead to any significant changes (FDR ≤ 0.05) in polysomal mRNA of parental HCT116 cells, and only 6 mRNA were changed in the total RNA sample (**Figure 3C, EV4D**). Similarly, the eIF3b-mAID cell line showed a small number of changes in polysomal mRNAs (22), an observation that is consistent with the unchanged P/M ratio after 12 h of IAA administration (**Figure 3B, EV5A**). 152 mRNAs were changed in the total mRNA sample indicating a distinct transcriptional response to short-term eIF3b depletion (**Figure EV4D**).

In contrast, depletion of eIF3a for 12 h led to a pronounced downregulation of 5049 polysomal mRNAs (**Figure 3C, EV4D**). Only 1.58% of these changes also registered in the total mRNA sample, indicating that the vast majority of the changes occurred at the translational level. Depletion of eIF3a also led to the induction of 328 mRNAs in the total mRNA sample but only 26 (7.9 %) of these mRNAs were significantly increased in the polysomal mRNA sample (**Figure EV4D**). This indicates that cells acutely depleted of eIF3a mount a transcriptional response, which fails to substantially perpetuate into protein production after 12 h of eIF3a depletion.

Cells depleted of eIF3e or eIF3f showed similar albeit blunted responses. 2561 mRNAs were downregulated in polysomes of eIF3e depleted cells of which 5% were also downregulated in the total mRNA sample. In eIF3f depleted cells, 1058 mRNAs were reduced in the polysomal fraction and 1.7% of these changes were reflected in the total mRNA sample (**Figure 3C, EV4D**). These differences in numbers are consistent with the less pronounced growth phenotypes of cells depleted of eIF3f compared to eIF3e (**Figure 2A, C**). In stark contrast, in eIF3k depleted cells, only 8 mRNAs were downregulated in polysomes, whereas 156 mRNAs were upregulated and only 3.2% of these changes occurred in the total mRNA sample (**Figure 3C, EV4D**).

Interrogating the polysomal changes for overrepresented Gene Ontology terms, no functional enrichments were obtained for eIF3a depleted cells. This indicates that eIF3a promotes the translation of a wide array of mRNAs without any detectable selectivity. Likewise, there was no enrichment for eIF3b depleted cells due to the low number of mRNAs affected 12 h after IAA. In contrast, depletion of eIF3e and eIF3f led to enrichment of an overlapping set of functional categories, including ribosomes, mitochondrial organization, and membrane processes in the sets of mRNAs whose polysomal association was decreased (**Figure 3D, Supplementary Data File 3**). These categories agree with those identified in our previous studies in yeast and MCF10A cells (Lin et al., 2020; Shah et al., 2016). In contrast, the GO terms “ribosomal subunit” and “polysome” was enriched in the set of 155 mRNA which increased in the polysomal fraction upon depletion of eIF3k, suggesting a role of eIF3k in regulating ribosome biosynthesis (**Figure 3D, Supplementary Data File 3**).

We also calculated translational efficiencies (TEs) defined as Log2 polysomal mRNA / total mRNA (**Supplementary Data File 4)**. As expected, median TEs were centered closely around 0 for parental HCT116 and eIF3b-mAID cells after 12 h of IAA (**Figure 3E**). TE fold change medians were strongly reduced for eIF3a (Log2_TE_ = -1.07), eIF3e (Log2_TE_ = -0.43) and eIF3f (Log2_TE_ = -0.28) depleted cells but increased for eIF3k depleted cells (Log2_TE_ = 0.15) (**Figure 3E**). The number of mRNAs whose TE changed (defined as changes at FDR ≤ 0.2) in the various cell lines correlated with the overall effect size on mRNA translation and ranged from 4951 mRNAs in eIF3a depleted cells to 1930, 103, and 0 in eIF3e, eIF3f and eIF3k depleted cells, respectively. The low number of TE changes in eIF3f and eIF3k depleted cells are due to the comparatively small effect size on translation which we could not measure reliably.

A potential limitation of extrapolating mRNA translation from the RNA-seq data is that co-fractionation with polyribosomes does not necessarily indicate that an mRNA is actively translated. We therefore performed an orthogonal analysis of protein synthesis by metabolic pulse labeling with heavy amino acids and protein identification by LC-MS/MS (pSILAC, (Selbach et al., 2008)). Six hours after the addition of IAA or the vehicle DMSO, eIF3-mAID cells were labeled with heavy (^13^C ^15^N) lysine and arginine for another 6 hours and, as with the RNA-seq experiments, cells were harvested for LC-MS/MS after a total of 12 h in IAA. Log2 fold changes (Heavy IAA/Heavy DMSO) in the labeling of eIF3 proficient and depleted cells with heavy amino acids were determined to assess eIF3 subunit-dependent differences in protein synthesis (**Supplementary Data File 5)**. Between 990 and 1771 proteins were measured in the various cell lines. Data for 1543 proteins from parental HCT116 cells showed that protein synthesis was not perturbed by IAA treatment with not a single protein differing at the FDR cutoff of 0.1 and the Log2 median fold change (Log2_pSILAC_) centered closely around 0 (**Figure 3F**).

In stark contrast, the synthesis of 811 out of the total of 1027 proteins measured (79 %) showed a reduction in eIF3a depleted cells for a median Log2 fold change of -3.16 (**Figure 3F**). Similar, albeit weaker reductions were measured upon depletion of eIF3e (median Log2_pSILAC_ = -0.52, 22.5% of measured proteins synthesis rates decreasing) and eIF3f (median Log2_pSILAC_ = -0.37, 2.1% of synthesis rates decreasing) (**Figure 3F**).

Surprisingly, protein synthesis was also downregulated in eIF3b-mAID cells 12 h after addition of IAA (median Log2_pSILAC_ = -0.72, **Figures 3F**) even so polysome profiles and translation efficiency (median Log2_TE_ = 0.0041) were unchanged at the same time point (**Figure 3B, EV5A)**. Among the 81 mRNAs downregulated in the total but not the polysomal mRNA sample upon treating eIF3b-mAID cells with IAA for 12 h, we noticed a strong enrichment of the GO terms “cellular amino acid metabolic process” and “organic acid transport” (**Figure EV5B**). Manual inspection revealed the downregulation of several mRNAs encoding amino acid transporters. These included SLC7A5 and SLC38A2 transporting neutral amino acids as well as SLC7A1 mediating the import of positively charged lysine and arginine which were used for stable isotope labeling in the pSILAC experiment. We confirmed 13 – 30% downregulation of these amino acid transporter mRNAs in total mRNA by qPCR but not in polysomal mRNA fractions (**Figure EV5C, D**). Thus, apparent downregulation of mRNAs encoding amino acid transporters at the transcriptional level is most likely responsible for the reduction in protein synthesis observed upon shut-off of eIF3b protein expression for 12 hours.

In agreement with the RNAseq-based TE, protein synthesis was increased in eIF3k depleted cells rather than decreased (median Log2_pSILAC_ = 1.23, 26% of synthesis rates increased, **Figure 3F**). Global agreement between RNA-seq (TE) and protein synthesis (pSILAC) data moving into the same directions was obtained for all eIF3 conditional cell lines with the exception of eIF3b-mAID as expected (**Figure EV5E**).

### eIF3k negatively modulates global protein synthesis through suppressing ribosomal protein synthesis

The most striking contrast between eIF3 subunits was the unique upregulation of cell and tumor growth and global translation in eIF3k depleted cells (**Figure 2, 3**), whereas cells depleted of all other eIF3 subunits showed the opposite behavior. We first considered the possibility that depletion of eIF3k causes increased growth by sensitizing cells to mitogenic signaling. To test this, we asked whether the growth advantage of eIF3k depleted cells would be enhanced at reduced concentrations of fetal bovine serum (FBS) thus indicating increased mitogen sensitivity. Reducing FBS from 10% to 5% or 2% did not lead to a greater advantage of eIF3k depleted cells (**Figure 4A**). Instead, eIF3k replete and depleted cells maintained their characteristic difference in cell number increase over time which was also reflected in the metabolic assay measuring NAD(P)H levels (CCK-8, **Figure EV6A**). We also did not observe any differences in serum stimulated mitogenic signaling with mTOR, S6 kinase, and Erk1/2 phosphorylation being induced with similar kinetics in eIF3k replete and depleted cells (**Figure 4B, EV6B**), findings that are consistent with the cell cycle kinetics in **Figure 2C**. Likewise, cells showed no difference in sensitivity to the mTOR inhibitor rapamycin depending on eIF3k status as determined by immunoblotting with phosphospecific mTOR antibodies (**Figure 4C, EV6C**). We did note a distinct increase in phosphorylated S6 kinase in eIF3k-depleted cells (**Figure 4C, EV6C**), suggesting that ribosome activity may be increased. In summary, these data suggest that increased growth of eIF3k depleted cells is not due to increased responsiveness to mitogenic stimulation.

**Figure 4.**
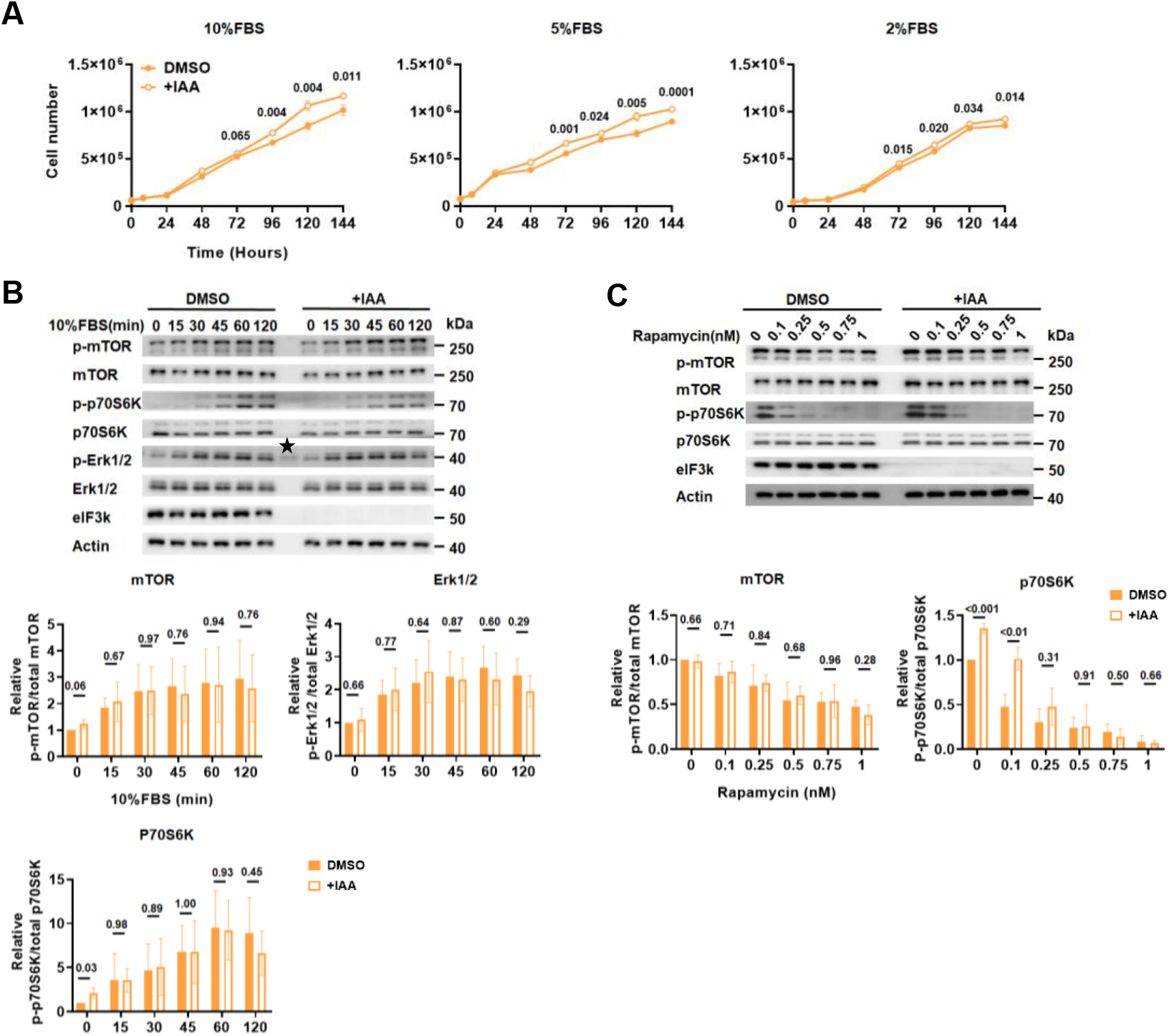
Effect of eIF3k depletion on growth factor sensitivity. **(A)** eIF3k-mAID cells were maintained in media containing different concentrations of FBS and DMSO or IAA as indicated, and cell numbers were determined at various time points. Data represent means ± SD, n = 3. Numbers indicate p values (two-stage step-up method of Benjamini, Krieger, and Yekutieli). **(B)** eIF3k-mAID cells were serum starved by maintaining in media containing 0% FBS for 24 h. During the last 12 h of starvation, DMSO or IAA was added as indicated. Cells were re-stimulated with media containing 10% FBS, and the expression of the indicated proteins was followed of a period of 120 minutes. The data from triplicate experiments were quantified and plotted as relative ratios of phosphorylated to unphosphorylated species. Bars represent means ± SD, n = 3 (see Figure S6B, C). Numbers indicate p values (unpaired Student’s t-test). **(C)** eIF3k-mAID cells were exposed to IAA or DMSO for 12 hours, followed by the addition of increasing concentrations of rapamycin for 1 h. The expression of the indicated proteins was determined by immunoblotting. The data from triplicate experiments were quantified and plotted as relative ratios of phosphorylated to unphosphorylated species. Bars represent means ± SD, n = 3 (see Figure S6B, C). Numbers indicate p values (unpaired Student’s t-test).

Since ribosome capacity is limiting for growth (Dai and Zhu, 2020), increased ribosome synthesis as apparent from the GO term enrichment (**Figure 3D**) might explain enhanced cell and tumor growth of eIF3k depleted cells. Indeed, as judged by sucrose density gradient profiling of lysate prepared from an equal number of cells, depletion of eIF3k for 12 or 48 h led to a 6.5 - 6.9% (p < 0.005) increase in total cellular ribosome content (**Figure 5A, Figure EV6D**). In line with the GO term enrichment analysis, inspection of the global transcriptomic and proteomic eIF3k-mAID datasets revealed a list of 12 ribosomal proteins (RPs) whose mRNAs were upregulated in polysomes but not in the total mRNA thus having an increased TE (**Supplementary Data File 6**). The corresponding proteins also showed increased synthesis by pSILAC (**Supplementary Data File 6**). We chose 3 of these mRNAs, *RPS15A (uS8), RPS4X (eS4)*, and *RPL7A (eL8)* to determine their partitioning into monosomal and polysomal mRNA fractions by RT-qPCR. Triplicate experiments confirmed increased polysomal association of RP encoding mRNAs 12 hours after adding IAA to eIF3k-mAID cells, while no such increase was seen for eIF3k-independent *PATZ1, PTPRU, JUNB*, and *HSPH1* mRNAs (**Figure 5B**). Similar results were obtained after extending eIF3k depletion to 48 hours (**Figure EV6E**). The data strongly suggest that RP mRNAs are targets of negative translational control by eIF3k.

**Figure 5.**
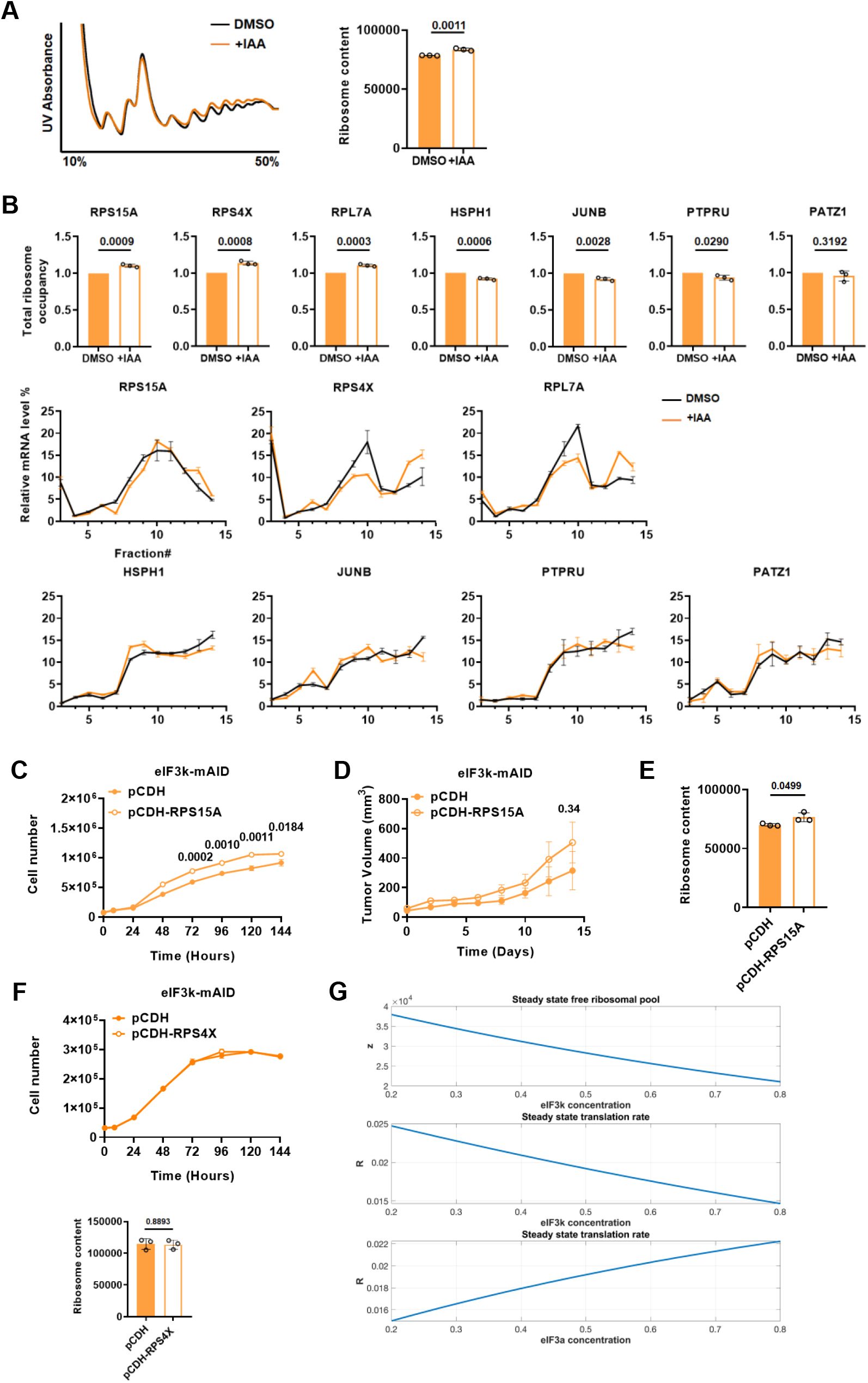
Effect of eIF3k depletion on ribosome content and RPS15A. **(A)** Lysates of eIF3k-mAID cells exposed to DMSO or IAA for 12 h were separated by sucrose density gradient centrifugation. Total ribosome content was determined by integrating the monosomal and polysomal peaks and plotted. Error bars represent means ± SD, n = 3. Numbers indicate p values (unpaired Student’s t-test). **(B)** Total ribosome occupancy of the indicated mRNAs in eIF3k-mAID cells exposed to DMSO or IAA for 12 hours was determined by RT-qPCR of RNA across a sucrose density gradient (see Materials and Methods). Bars represent means ± SD, n = 3; numbers indicate p values (unpaired Student’s t-test). Triplicate RT-qPCR data across the sucrose gradient are shown below the bar graphs. **(C)** Equal numbers of eIF3k-mAID cells stably expressing ectopic RPS15A (pCDH-RPS15A) or empty vector (pCDH) were plated and counted over a period of 6 days. Graphs represent means ± SD, n = 3. Numbers indicate p values (two-stage step-up method of Benjamini, Krieger, and Yekutieli). **(D)** 1 × 10^6^ eIF3k-mAID cells stably expressing ectopic RPS15A (pCDH-RPS15A) or empty vector (pCDH) were injected into nude mice and tumor growth was followed for 2 weeks. Graphs represent means ± SD, n = 5 to 7 (see Figure S6F). Numbers indicate p values (two-stage step-up method of Benjamini, Krieger, and Yekutieli). **(E)** Lysate of eIF3k-mAID cells stably expressing ectopic RPS15A (pCDH-RPS15A) or empty vector (pCDH) were separated by sucrose density gradient centrifugation. Total ribosome content was determined by integrating and summing the monosomal and polysomal peak areas. Error bars represent means ± SD, n = 3. Numbers indicate p values (unpaired Student’s t-test). **(F)** Equal numbers of eIF3k-mAID cells stably expressing ectopic RPS4X (pCDH-RPS4X) or empty vector (pCDH) were plated and counted over a period of 6 days. Graphs represent means ± SD, n = 3. Numbers indicate p values (two-stage step-up method of Benjamini, Krieger, and Yekutieli). Total ribosome content of these cells was determined as described in (A). **(G)** Steady-state free ribosomal pool and translation rate as a function of eIF3k concentration (top two graphs) and the steady-state translation rate as a function of eIF3a concentration (bottom graph). The concentration of other eIF3 subunits is kept constant at 0.5.

RPS15A is special because it is frequently overexpressed in human colorectal cancers (Chen et al., 2016), and its overexpression was shown to be sufficient to promote tumor growth in mice, although its effects on translation were not assessed (Guo et al., 2018; Liu et al., 2019). We found that stable overexpression of *RPS15A* mimicked the effect of eIF3k depletion on cell growth, leading to a marked stimulation (**Figure 5C**). Likewise, *RPS15A* overexpression was sufficient to increase tumor growth (**Figure 5D, EV6F**) and the total cellular ribosome content (**Figure 5E**). No stimulation of cell growth and ribosome content was obtained upon ectopic expression of RPS4X (**Figure 5F**). These data suggest that depletion of eIF3k promotes cell and tumor growth by increasing cellular ribosome content.

To gauge the plausibility of this conjecture, we sought to integrate our experimental data into a mathematical-computational predictive model. We adopted our previously developed ribosome flow model (Reuveni et al., 2011; Zarai and Tuller, 2018) that captures fundamental aspects of translation, including: (i.) Potential differences in decoding-time for each codon, which is related to the local biophysical properties of the mRNA (e.g., its folding and the encoded amino acids) and its interaction with translation factors and/or the availability of such factors (e.g., tRNA levels); (ii.) Initiation rates, which are affected by the properties of the mRNA and initiation factors as well as global factors such as the concentrations of ribosomes and translation factors; (ii.) Multiple ribosomes translating the same mRNA at a certain point in time; (iv.) Excluded volume interactions between ribosomes and possible traffic jams; (v.) Directionality of the ribosome flow (the flow is totally asymmetric. i.e. unidirectional); (vi.) The fact that mRNA molecules compete for a finite pool of ribosomes. Based on parameters derived from ribosome foot printing data of HEK293 cells (Dana and Tuller, 2015), the model can predict the exact relations between the levels of distinct eIF3 subunits and global translation variables such as translation rates and the free ribosomal pool which are expected to be positively correlated with growth rate. Using TP53 as a model mRNA, we simulated the effect of the eIF3k concentration on the free ribosomal pool and the translation rate, as well as the effect of the eIF3a concentration on the translation rate. As apparent in **Figure 5G**, decreasing the eIF3k concentration increases the steady state free ribosomal pool which in turn increases the translation rate. In contrast, decreasing eIF3a concentration decreases the steady state translation rate via its effect on initiation. Similar results were obtained for simulating steady-state average ribosome density (ρ) and steady-state average ribosome jams (θ) (**Figure EV6G**). Thus, the simulations readily capture key experimental observations of our study: (i.) eIF3k has a negative effect on the ribosome pool, and thus on the global initiation rate, translation rate, and growth. eIF3a positively affects the translation rate via initiation.

### eIF3k-dependent suppression of *RPS15A* mRNA translation is mediated by eIF3 binding to the 5’-UTR

Our results suggested that eIF3k is a negative modulator of the translation of RP mRNAs, especially *RPS15A*. To address mechanisms of this regulation, we examined global CLIP datasets (Lee et al., 2015; Meyer et al., 2015) for the presence of eIF3 binding sites in each of the 12 RP mRNAs we found translationally upregulated in eIF3k depleted cells. Only two mRNAs, those encoding RPS15A and RPS4X, were reported to have eIF3 binding sites identified in both CLIP datasets (**Figure 6A**). To assess the requirement of these binding sites for the translation of *RPS15A* and *RPS4X* mRNAs, we performed CRISPR/Cas9 genome editing to delete the binding sites from both alleles of eIF3k-mAID cells. We could not obtain viable cells upon editing *RPS4X*. We subsequently determined by long-read RNA sequencing (data not shown) that the mapped eIF3 binding site is only present in a minor isoform of *RPS4X* mRNA (∼5% of the total pool in human lymphocytes, **Figure 6A**). Genomic deletion of the eIF3 binding site is predicted to disrupt the promoter of the major mRNA species thus likely abolishing the transcription of ∼95% of the *RPS4X* mRNA and leading to lethality. This does not rule out a role of the eIF3 binding site in increasing the translation of the minor isoform of *RPS4X* mRNA in eIF3k depleted cells, but this cannot be addressed by genomic deletion of the mapped binding site.

**Figure 6.**
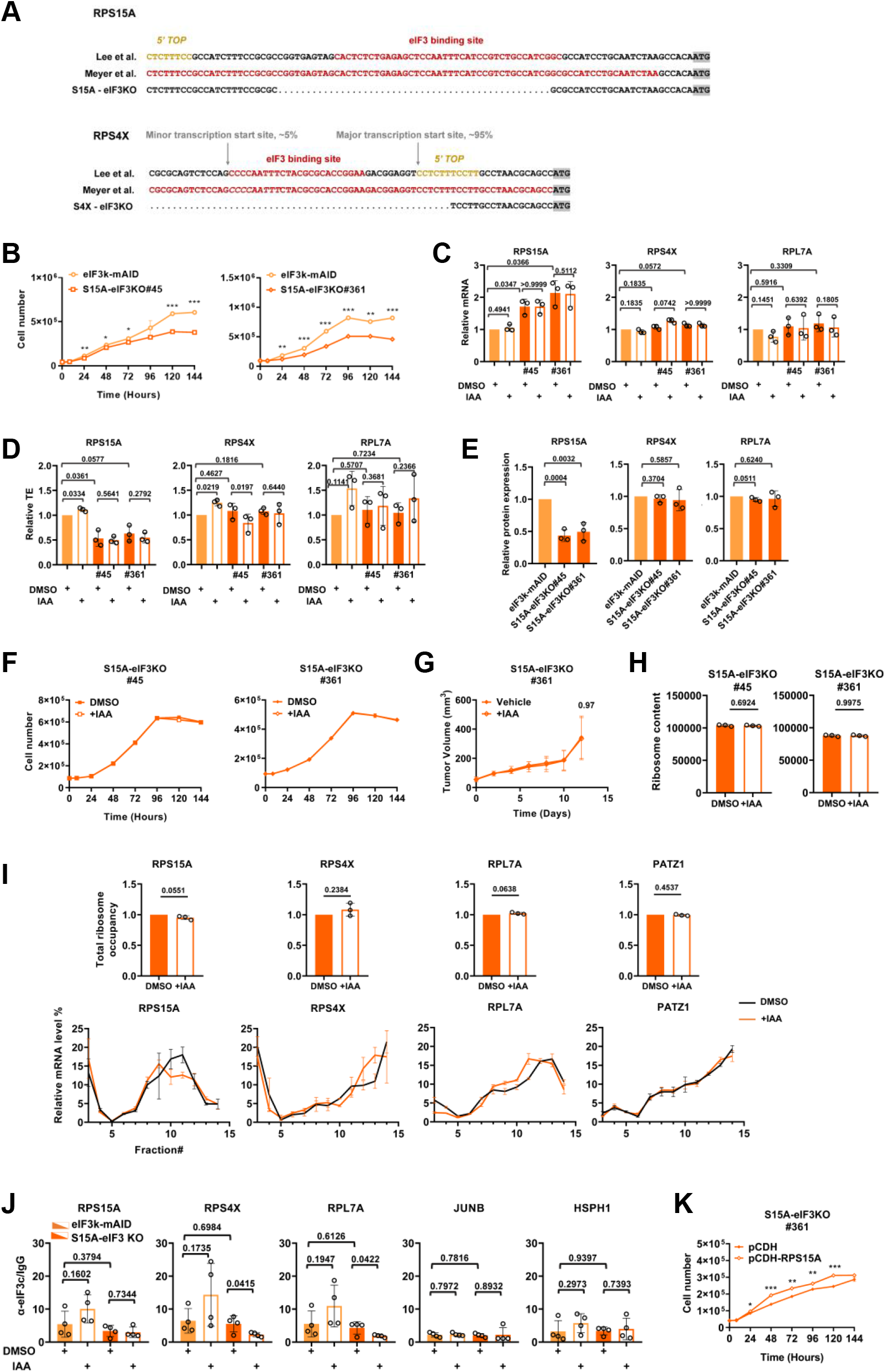
Effect of the eIF3 binding site on cell proliferation, tumor growth, and RPS15A. **(A)** 5’-UTR sequences of RPS15A and RPS4X. 5’-TOP element known to boost translation of ribosomal protein mRNAs (Meyuhas, 2000) are highlighted in gold prints, the eIF3 binding sites mapped by Lee et al. (Lee et al., 2015) and Meyer et al. (Meyer et al., 2015) are highlighted in red print. The alleles created by gene editing are shown (S15A-eIF3KO, S4X-eIF3KO). Note that the eIF3 binding site in RPS4X is upstream of or overlapping with the major transcription start site. Thus, the S4X-eIF3KO truncation we failed to make most likely abolished the transcription of RPS4X mRNA. **(B)** Parental eIF3k-mAID cells or S15A-eIF3KO cells (clones #45 and # 361) were maintained in standard media, and cell numbers were determined at various time points. Data represent means ± SD, n = 3. Asterisks denote: * p < 0.005, ** p < 0.00005, *** p < 0.000005 (two-stage step-up method of Benjamini, Krieger, and Yekutieli). **(C)** Relative levels of mRNAs encoding *RPS15A, RPS4X*, and *RPL7A* before and after depletion of eIF3k were determined in the indicated cell lines by RT-qPCR. Data were normalized to the signal obtained for GAPDH. Bars represent means ± SD, n = 3; numbers indicate p values (unpaired Student’s t-test). **(D)** Relative translational efficiencies (TEs) of mRNAs encoding *RPS15A, RPS4X*, and *RPL7A* before and after depletion of eIF3k were determined in the indicated cell lines. RT-qPCR was performed on total RNA and on RNA contained within polysomal fractions > 2 ribosomes, and TE was calculated according to the formula TE = polysomal mRNA / total mRNA. Bars represent means ± SD, n = 3; numbers indicate p values (unpaired Student’s t-test). **(E)** Basal expression of the indicated proteins was determined in parental eIF3k-mAID cells or S15A-eIF3KO cells (clones #45 and # 361) by immunoblotting, followed by quantification of the blots. Bars represent means ± SD, n = 3 (Figure S7A). Numbers indicate p values (unpaired Student’s t-test). **(F)** Parental eIF3k-mAID cells or S15A-eIF3KO cells (clones #45 and # 361) were maintained in media with DMSO or IAA, and cell numbers were determined at various time points. Data represent means ± SD (too small to be visible), n = 3. All p values (two-stage step-up method of Benjamini, Krieger, and Yekutieli) were > 0.45 except at the single time point where indicated otherwise. **(G)** 1 × 10^6^ S15A-eIF3KO (clone #361) cells were injected into nude mice. When tumors reached a diameter of ∼ 4 mm, mice were treated with vehicle or 500 mg/kg IAA, and tumor growth was measured for 2 weeks. Graphs represent means ± SD, n = 5 to 7. Numbers indicate p values (two-stage step-up method of Benjamini, Krieger, and Yekutieli). **(H)** Lysate of S15A-eIF3KO cells (clones #45 and # 361) were separated by sucrose density gradient centrifugation. Total ribosome content was determined by integrating and summing the monosomal and polysomal peak areas. Error bars represent means ± SD, n = 3 (see Figure S7B). Numbers indicate p values (unpaired Student’s t-test). **(I)** Total ribosome occupancy of the indicated mRNAs in S15A-eIF3KO (clone #361) cells exposed to DMSO or IAA for 12 hours was determined by RT-qPCR of RNA across a sucrose density gradient (see Materials and Methods). Bars represent means ± SD, n = 3; numbers indicate p values (unpaired Student’s t-test). Triplicate RT-qPCR data across the sucrose gradient are shown below the bar graphs. **(J)** RNA immunoprecipitation. eIF3k-mAID and S15A-eIF3KO (clone #361) cells were exposed to DMSO or IAA for 12 hours. Cell lysates were employed in immunoprecipitation with eIF3c antibodies and co-precipitated mRNAs were quantified by qPCR. Bars represent means ± SD, n = 4; numbers indicate p values (unpaired Student’s t-test). **(K)** Equal numbers of S15A-eIF3KO (clone #361) cells stably expressing ectopic RPS15A (pCDH-RPS15A) or empty vector (pCDH) were plated and counted over a period of 6 days. Graphs represent means ± SD, n = 3. Asterisks denote: * p < 0.005, ** p < 0.00005, *** p < 0.000005 (two-stage step-up method of Benjamini, Krieger, and Yekutieli).

RPS15A 5’-UTR edited cell lines were obtained, however, which were denoted S15A-eIF3KO clones #45 and #361. While the edited cells grew slower than the parental eIF3k-mAID cells (**Figure 6B**), they had ∼2-fold increased *RPS15A* but not *RPS4X* or *RPL7* mRNA levels (**Figure 6C**). However, TE of *RPS15A* mRNA was ∼2-fold lower in the 5’-UTR edited cell lines (**Figure 6D**), observations that were consistent with a 2-fold reduction in RPS15A but not RPS4X or RPL7 protein (**Figure 6E, EV7A**). These data indicate that the eIF3 binding site in the 5’-UTR of the *RPS15A* mRNA acts as an amplifier of *RPS15A* mRNA translation. eIF3k and eIF3l were also reduced in S15A-eIF3KO cells, possibly as a compensatory measure aimed at boosting *RPS15A* translation (**Figure EV7A**).

Despite these baseline changes, S15A-eIF3KO cells depleted for eIF3k did not show the increase in growth rate found in the parental eIF3k-mAID cells (**Figure 6F**). Likewise, the growth of tumors derived from S15A-eIF3KO cells was not stimulated upon eIF3k depletion (**Figure 6G**). In agreement with these growth phenotypes, eIF3k depleted S15A-eIF3KO cells did not shown an increase in total ribosome content (**Figure 6H, EV7B**). Unlike in parental eIF3k-mAID cells, depletion of eIF3k in S15A-eIF3KO cells did not lead to increased recruitment of *RPS15A* mRNA into polysomes, whereas the stimulation was largely maintained for *RPS4X* and *RPL7A* mRNAs (**Figure 6I, EV7C**). Likewise, the increase in the TE of *RPS15A* mRNA observed in eIF3k-depleted parental cells was not apparent in 5’ -UTR edited S15A-eIF3KO cells (**Figure 6D**). Finally, RNA immunoprecipitation with eIF3c antibodies showed that binding of eIF3 to *RPS15A* mRNA was diminished in S15A-eIF3KO cells but increased in eIF3k-mAID cells upon downregulation of eIF3k (**Figure 6J**). Importantly, ectopic expression of RPS15A rescued proliferation of S15A-eIF3KO cells (**Figure 6K**), indicating that failure to augment RPS15A levels is the primary reason for the lack of growth stimulation of S15A-eIF3KO cells by depletion of eIF3k. In summary, these data suggest that physical interaction of eIF3 with the 5’-UTR of *RPS15A* imparts critical control on *RPS15A* mRNA translation and tumor cell growth.

### Depletion of eIF3k confers stress resistance dependent on the eIF3 binding site in *RPS15A* mRNA

Deletion of eIF3k and its binding partner eIF3l was previously shown to confer resistance to tunicamycin-induced endoplasmic reticulum (ER) stress, although no mechanistic explanation was provided (Cattie et al., 2016). Testing eIF3-mAID cell lines for sensitivity to 2 µg/ml tunicamycin, we found that depletion of eIF3k conferred resistance, whereas cell lines depleted of any of the other eIF3 subunits were sensitive in the CCK-8 assay (**Figure 7A, EV7D**). eIF3k-mAID cells ectopically expressing RPS15A showed the same resistance as cells depleted of eIF3k, whereas eIF3k depletion did not affect the sensitivity of S15A-eIF3KO cells (**Figure 7A, EV7D**). Similar behaviors of the various cell lines were observed for oxidative stress induced by tert-butyl hydroperoxide (**Figure 7B, EV7E**). Taken together, these results link eIF3k-dependent stress resistance to its role in controlling *RPS15A* mRNA translation.

**Figure 7.**
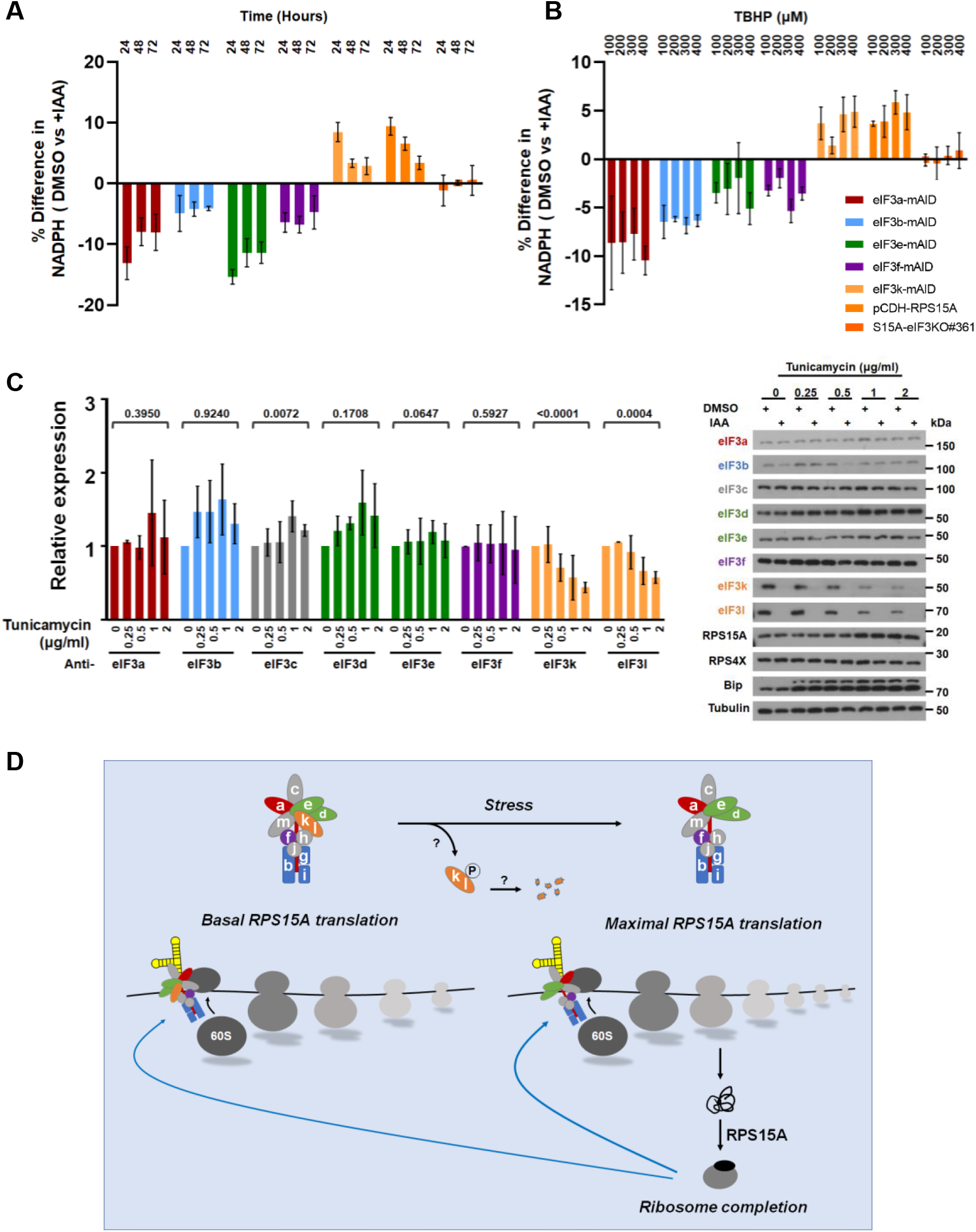
Effect of eIF3k on stress sensitivity. **(A)** The indicated eIF3-mAID cell lines were treated with DMSO or IAA for 12 hours, followed by exposure to 2 µg/ml tunicamycin for up to 72 hours. Metabolic activity (i.e. NAD(P)H) as a proxy of cell viability was determined by CCK8 assay. Graphs represent the percentage of change in cell viability upon downregulating eIF3 subunits with IAA. Date are means ± SD, n = 3. **(B)** Same experiment as in (A) but cells were exposed to increasing concentrations of the oxidative stress inducer tert-butyl hydroperoxide (TBHP) for 2 hours. **(C)** The indicated eIF3-mAID cell lines were treated with DMSO or IAA for 12 hours, followed by exposure to 2 µg/ml tunicamycin for 24 hours. The expression of individual eIF3 subunits, ribosomal proteins, and the ER stress marker BIP were assessed by immunoblotting. Tubulin is show for reference. The data from triplicate experiments were quantified and to avoid overcrowding of the graph data obtained from IAA-treated cells were not plotted. Bars represent means ± SD, n = 3 (see Figure S7F). Numbers indicate p values (unpaired Student’s t-test). **(D)** Model of the dual role of eIF3k in the translation of *RPS15A* mRNA. When the eIF3 binding site in the 5’-UTR is occupied by holo-eIF3 containing the eIF3k-l module, *RPS15A* mRNA is translated efficiently at basal rate. When the eIF3k-l module is missing (or functionally inactivated, perhaps degraded in response to stress conditions), the remaining eIF3 complex boosts maximal translation of *RPS15A* mRNA through the eIF3 binding site. The extra RPS15A then completes substoichiometric ribosomes to boost ribosome content, translation and growth.

Finally, we determined the effect of tunicamycin on the levels of individual eIF3 subunits. Whereas 2 µg/ml tunicamycin for 24 hours did not substantially affect the levels of eIF3a, b, c, d, e, and f, eIF3k and eIF3l were selectively downregulated by the treatment (**Figure 7C, EV7F**). Whereas the underlying mechanisms remain to be established, this response provides a facile mode for creating a cellular eIF3 complex devoid of the eIF3k-l module that could boost cell fitness and tumor growth through RPS15A-dependent increases in ribosome content.

## DISCUSSION

### Modular architecture of eIF3 and mRNA selectivity

Our quantitative data on eIF3 complex formation and mRNA translation in cells acutely depleted of eIF3 subunits reinforce our model that holo-eIF3 consists of functionally distinct modules with distinct mRNA selectivity (Lin et al., 2020; Shah et al., 2016). Our data are consistent with previous studies relying on knockdown of eIF3 subunits for 72 hours (Lin et al., 2020; Wagner et al., 2014, 2016), but due to the acute depletion afforded by the mAID system, we can assess the effects of eIF3 subunit depletion on holo-eIF3 and mRNA translation largely independent of compensatory effects in response to long-term subunit depletion.

Consistent with its role as a nucleation core of holo-eIF3 formation (Smith et al., 2016; Wagner et al., 2016), eIF3a was found to be a global regulator of mRNA translation affecting upwards of 5,000 mRNAs in HCT116 cells. The impact of eIF3a is so pervasive that no functional categories were enriched in its set of target mRNAs. Since depletion of eIF3a leads to disassembly of the octameric core complex, it is likely that it abolishes all functions of holo-eIF3 in mRNA translation.

eIF3e affects a far more restricted set of mRNAs, many encoding proteins engaged in mitochondrial and membrane-associated functions. This confirmed our previous findings in fission yeast and human MCF10A cells where eIF3e - likely in complex with eIF3d - functions to promote early translation elongation of mRNAs encoding membrane-associated proteins (Lin et al., 2020; Shah et al., 2016). Short-term depletion of eIF3f had a similar effect on holo-eIF3 as depletion of eIF3e and regulated an overlapping set of mRNAs, although its effects on mRNA translation and cell proliferation were less pronounced. This is surprising considering that eIF3f but not eIF3e is essential for viability in S. pombe (Zhou et al., 2005). Regardless, eIF3f target mRNAs enriched similar mitochondrial and membrane-associated functional categories as the eIF3e dataset, although it is presently unclear whether eIF3f also affects early translation elongation. Our data from HCT116 cells do not support previous conclusions that eIF3f is an inhibitor of global translation (Shi et al., 2006); only 17 mRNAs were increased in the polysomal fraction upon depletion of eIF3f (**Figure EV4D**).

Surprisingly, short-term downregulation of eIF3b by ∼73% did not affect mRNA translation despite beginning depletion of the eIF3b-g-i subcomplex. Even at strongly reduced eIF3b levels, cells maintained amounts of holo-eIF3 that are ostensibly sufficient for global mRNA translation (**Figure 1C**). Nevertheless, as shown by pSILAC proteomics, protein synthesis was reduced in cells depleted of eIF3b for 12 h. This reduction correlated with decreased levels of mRNAs encoding amino acid transporters and tRNA synthetases (**Figure EV5B, C**), pointing to a possible moonlighting function of eIF3b in mRNA transcription or stability control that may be more sensitive to acute changes in eIF3b levels than its function in translation. Consistent with this possibility, changes at the level of total mRNA are most abundant in eIF3b depleted cells compared to all other eIF3-mAID cell lines (**Figure EV4D**) and a fraction of eIF3b is known to localize to the cell nucleus (https://www.proteinatlas.org/ENSG00000106263-EIF3B/subcellular#human).

Despite quantitative and qualitative differences in mRNA selectivity, all tested eIF3 subunits except eIF3k were required for cell and tumor growth. This is consistent with cell autonomous pro-oncogenic functions of these eIF3 subunits and renders studies suggesting tumor suppressor functions of eIF3e and eIF3f difficult to reconcile (Marchione et al., 2013; Sesen et al., 2017). It is noteworthy in this context that eIF3f is essential in fission yeast (Zhou et al., 2005) and eIF3f and eIF3e homozygous knockout mice are early embryonic lethal (Docquier et al., 2019; Lin et al., 2020; Sadato et al., 2018). It is thus unclear what tumor cells stand to gain from reducing the levels of these essential growth promoting factors.

### Dual roles of eIF3 in the regulation of *RPS15A* mRNA translation: eIF3k as a regulator of ribosome reserve capacity

eIF3k was found to be unique in that its depletion promoted cell and tumor growth, effects presumably mediated by the eIF3k/l module. This unusual gain of fitness was not attributable to increased sensitivity to mitogenic signaling but rather to an increase in total ribosome content and global translation. The observed inverse relationships between eIF3k levels with ribosome content and global translation were recapitulated in the RFM model of mRNA translation thus underpinning our experimental observations by computational modeling.

Among several RP mRNAs that are translationally upregulated in eIF3k depleted cells, only *RPS15A* is known to bind eIF3 in its 5’-UTR (Lee et al., 2015; Meyer et al., 2015). Deleting this binding site had two effects: (i.) It reduced cell proliferation and caused a ∼50% decrease in RPS15A protein. (ii.) It prevented the increase in ribosome content and cell and tumor growth induced by depletion of eIF3k. Thus, the eIF3 binding site has a dual function in controlling RPS15A: (i.) It positively regulates the basal translation of *RPS15A* mRNA, and (ii.) it negatively regulates the maximal rate of *RPS15A* mRNA translation via eIF3k (**Figure 7D**). A repressive effect of eIF3 via 5’-UTR binding has previously been described for the mRNA encoding ferritin light chain (Pulos-Holmes et al., 2019), although the repression was not attributed to a distinct subunit. eIF3k-mediated repression of *RPS15A* translation thus represents – to our knowledge - the first example of subunit-selective translational repression by eIF3 through an element in the 5’-UTR. Dual positive and negative functions of nucleic acid binding motifs depending on the subunit composition or posttranslational modification of binding protein complexes are well established in transcriptional regulation (Henley and Dick, 2012). In conclusion, through eIF3k, eIF3 functions as a rheostat to keep *RPS15A* mRNA translation within relatively narrow boundaries.

An intriguing question is how cellular ribosome content can be increased by boosting the translation of individual mRNAs encoding ribosomal proteins. Remarkably, whereas ectopic expression of several RPs induces nucleolar stress and p53-dependent growth arrest (Jin et al., 2004; Lohrum et al., 2003; Zhang et al., 2003), ectopic RPS15A increased ribosome content and augmented cell and tumor growth. This suggests that RPS15A can mobilize reserve ribosome capacity, possibly by completing a pool of sub-stoichiometric ribosomes known to exist in S. cerevisiae (Yu et al., 2020). Similarly, completion of ribosome subunit stoichiometry by boosting the expression of sub-stoichiometric ribosomal proteins was linked to increased cell growth (Slavov et al., 2015) and neuronal mRNA translation (Fusco et al., 2021). Whereas eIF3k depletion led to increased translation of at least 11 additional RPs (**Supplementary Data File 6**), an effect that was reflected in the enrichment of the GO term “polysomes” in our global translatomic datasets (**Figure 3D**), RPS15A appears uniquely required for increasing ribosome content and growth.

What might be the physiological role of the proposed function of eIF3k as a rheostat for keeping *RPS15A* mRNA translation within relatively narrow limits? The general picture emerging from studies in microbes (Korem Kohanim et al., 2018; Mori et al., 2017) and mammalian systems (Slavov et al., 2015) is that reserve ribosome capacity is maintained in a trade-off between maximum exponential growth rate versus the ability to respond quickly to changes in environment. For example, ribosome capacity can become limiting under environmental stress conditions when cells are swamped with stress-induced transcripts that need to be turned into new proteins quickly (Lee et al., 2011). Our data introduce eIF3k as a new regulator of ribosome capacity at the level of translation. A possible model is that stress signaling triggers the loss or functional inactivation of eIF3k and eIF3l such that the residual eIF3 complex lacking these subunits maximally stimulates *RPS15A* translation (**Figure 7D**). Consistent with this model is the observation that loss-of-function mutations in eIF3k and eIF3l lead to enhanced resistance of C. elegans to endoplasmic reticulum stress induced by tunicamycin (Cattie et al., 2016), a phenotype we reproduced and extended to oxidative stress in HCT116 cells (**Figure 7A, B**).

Within the tumor microenvironment, cancer cells are well known to be exposed to rapid fluctuations in nutrient, pH, and redox conditions (Luo et al., 2009). In light of our discoveries, the downregulation of eIF3k we identified in human colon and renal cancers may enable tumor cells to tap into ribosome reserves to maximize stress resilience without a substantial cost on growth. If so, this stress mitigation pathway may present an attractive target for interventions geared toward disabling eIF3-boosted ribosome capacity.

## MATERIALS and METHODS

### Cell lines

The male HCT116 cell line expressing OsTIR1 (Natsume et al., 2016) was obtained from the Riken Bioresource Research Center of Japan and was cultured in McCoy’s 5A medium (Thermo Fisher Scientific) supplemented with 10% FBS (Gibco), 2 mM L-glutamine,100 U/ml penicillin, and 100 mg/ml streptomycin at 37°C and 5% CO_2_. Cells were periodically determined to be free of mycoplasma.

### Mice

BALB/c male nude mice (4–6 weeks of age) were used for subcutaneous xenografts. Mice were housed under pathogen-free conditions and maintained on a 12 h light/12 h dark cycle with food and water supplied ad libitum.

### Generation of cell lines

To generate cell lines in which both alleles of eIF3 subunits were tagged with mAID-mClover and mAID-mCherry, donor plasmids were constructed according to (Natsume et al., 2016). Homology arm fragments for eIF3a, b, e, f, and k containing a central BamHI site were obtained by gene synthesis and cloned into a donor plasmid. In a second step, the respective mAID-mClover and mAID-mCherry cassettes containing a selection marker were cloned at the BamHI site between the homology arms. Homology arms were designed to mutate the recognition sequence after integration at the target locus to prevent re-cutting. HCT116-OsTIR1 cells were grown in 6-well plates before 800 ng Cas9 plasmid, and 600 ng mAID donor plasmids (**Reagent Table**) were transfected using 8 μl Lipo2000. 48h after transfection, cells were re-plated on 10 cm dishes, followed and selected with 700 μg/ml G418 and 100 μg/ml HygroGold. After 12 days, colonies were picked and verified by PCR and Sanger sequencing and by immunoblotting. The list of primers for various truncation and point mutants are summarized in the Key Resource Table. To induce the degradation of mAID-fused proteins, 500 µM indole-3-acetic acid (IAA) was added to the culture medium.

To generate eIF3k-mAID cells in which the eIF3 binding sites in the 5’-UTRs of RPS15A and RPS4X were deleted, targeting plasmids were constructed according to the protocol by Ran et al. (Ran et al., 2013). Cas9 cutting sites 3’ and 5’ of the region to be deleted were identified using a web tool (http://crispor.tefor.net/, (Concordet and Haeussler, 2018)). Pairs of corresponding sgRNAs were individually cloned into either BsmBI-digested lentiCRISPR v2-Blast or lentiCRISPR v2-Puro. To produce virus, HEK293T cells at ∼50% confluence were transfected in 35 mm dishes with 0.5 μg pMD2.G and 0.75 μg psPAX2 packaging vectors and 1 μg of the lentiCRISPR v2 plasmid with 5 μl Lipo2000 in optiMEM. The medium was exchanged for DMEM high glucose medium with 10% FBS after 6h. Virus supernatant was collected after 48h, passed through a 0.45 μm filter, and stored at −80°C. 30% confluent eIF3k-mAID cells were infected by pairs of sgRNA virus in 35 mm dishes with 8 μg/ml polybrene. After 24h, the medium was exchanged for McCoy’s 5A, and cells were selected for one week in puromycin (4 μg/ml) and blasticidin (30 μg/ml). A small fraction of the infected cells was collected to determine the efficiency of CRISPR editing. Different combinations of guide RNAs showed different efficiency. Cells infected with the most efficient pair of guide RNA viruses were expanded, and single colonies were picked and confirmed by genomic PCR and Sanger sequencing. The list of primers for various truncation are summarized in the **Table V1**.

For stable overexpression of RPS15A and RPS4X in eIF3k-mAID cells, a recombinant lentivirus was constructed by inserting the PCR-amplified cDNA sequence into XhoI and XbaI sites of pCDH-CMV-MCS-EF1-puro-3xFlag-3xHA (generously provided by L. Wen). 293T cells were transfected with the packaging plasmids pMDLg/pRRE, pVSV-G, and pRSV-Rev at a ratio of 0.5 : 0.3 : 0.2 μg and 1 μg pCDH-CMV-RPS15A-3xHA or pCDH-CMV-RPS4X-3xHA plasmids. Virus collection and infection was as described above. After infection, cells were selected with puromycin (4 μg/ml) for 7 days and lysate of polyclonal cultures were tested for RPS15A expression by immunoblotting.

### Proliferation/viability assays

Commonly used tetrazolium dye reduction assays measuring NAD(P)H levels (MTT and CCK-8) were employed as surrogate measures of cell proliferation/viability. 8 × 10^3^ cells per well were seeded into 96 well plates. Cells were incubated with 10 μl of CCK8 reagent and incubated at 37°C until color development. Absorbance measurements were performed at 450 nm using a microplate reader (Thermo Scientific™ Multiskan™ FC). For MTT assays, 20 μl of MTT reagent at 5 mg/ml were added into each well for a final concentration of 0.5% and incubated for 4 hours at 37°C. The medium was removed and 100 μl 100% DMSO was added to each well followed by vigorous mixing. After incubation at room temperature for 10 minutes, samples were mixed, and absorbance was measured at 540 nm in a microplate reader (Thermo Scientific™ Multiskan™ FC).

### Colony formation assay

1 × 10^3^ cells were plated on 100 mm culture dishes in 3 replicates and cultured for 2 weeks. After removal of media and washing in PBS, 10 ml methanol was added, and cells were fixed for 30 min. Upon removal of methanol, cells were stained with crystal violet in 20% methanol in PBS for 30 min, rinsed with water, and dried. Colony numbers were determined with CFU Scope (http://www.cfu.ai).

### Cell cycle analysis by flow cytometry

Cells were collected by trypsinization and washed with cold PBS and then fixed in cold 70% ethanol in 4°C for a minimum of 2h. Fixed cells were treated with 100 µg/mL RNase and stained with 50 µg/ml propidium iodide in PBS for 15 – 30 min in the dark. Samples were run on an ATTUNE NXT Flow Cytometer (Thermo Fisher Scientific) and data was analyzed with FlowJo software.

### Xenografts studies

Animal experiments were performed in accordance with the Guiding Principles in the Care and Use of Animals (China) and were approved by the Laboratory Animal Ethics Committee of Xiamen University. 1 × 10^6^ cells in a volume of 100 µl were inoculated subcutaneously into nude mice. After tumor reached a diameter of ∼4 mm (∼50 mm^3^), IAA (500 mg/kg body weight in corn oil) or an equal amount of corn oil vehicle was administered by oral gavage daily for two weeks. Tumor sizes were monitored every two days by measurement with a caliper and body weights were recorded. Tumor volumes were calculated according to the formula: volume = width^2^ × length/2. Mice were sacrificed after two weeks, and tumors were excised for analysis. Tumor imaging was done on a Caliper IVIS Lumina II instrument with imaging mode set to Fluorescence, exposure time to 1 min, and excitation and emission filter set to 465 nm and 515-575 nm, respectively.

### Immunoblotting

For protein extraction, cells were washed twice with cold PBS, scraped off and lysed in SDS sample buffer (60 mM Tris/HCl, pH 6.8, 5% beta-mercaptoethanol, 2% SDS, 10% glycerol, 0.02% bromophenol blue) followed by heating for 8 min at 95°C. Lysates were separated by SDS-PAGE and immunoblotting was done as described (Tian et al., 2021). Primary antibodies used in this study were listed in the **Reagent Table**. Quantification of band intensities was done using ImageJ software.

### Native PAGE analysis

Cells contained in a 35 mm dish at 70 - 80% confluence were lysed in 250 μl native lysis buffer supplemented with 1% digitonin by pipetting up and down 10 times and incubation on ice for 10 min. Lysates were centrifuged at 20,000 g for 30 min at 4°C. Protein concentration was measured by BCA protein assay and Coomassie blue G-250 was added to a final concentration of 0.25%. Equal amounts of protein (20 μg) were resolved on 4-16% Native PAGE gels. Electrophoresis and immunoblotting were done according to the manufacturer’s manual (Thermo Fisher).

### Sucrose density gradient separation of ribosomes and RT-qPCR

10 min prior to harvest, 100 μg/ml cycloheximide (CHX) was added to cells contained in two 15 cm dishes at approximately 80% confluence. Cells were incubated in 0.5 ml hypotonic buffer (5 mM Tris/HCl, pH 7.5, 2.5 mM MgCl_2_, 1.5 mM KCl and 1 x Pierce™ protease inhibitor cocktail) supplemented with 100 μg/ml CHX, 1 mM DTT, and 100 units of RNase inhibitor on ice for 20 min and vortexed for 15 s. Triton X-100 and sodium deoxycholate were added to a final concentration of 0.5%, and the lysate was vortexed for another 5 s. The cell lysates were centrifuged at 16,500 g, at 4°C for 7 min. Supernatants were collected and absorbance at 260 nm was measured. 20 - 30 OD_260_ of lysate was gently layered over 10– 50% sucrose gradients in buffer (20 mM HEPES-KOH, pH 7.6, 5 mM MgCl_2_, 100 mM KCl, 100 μg/ml CHX, 10 units/ml RNase inhibitor and 1 x Pierce™ protease inhibitor cocktail). Gradients were centrifuged at 36,000 rpm (Beckman, SW41Ti) for 2 h at 4°C. After centrifugation, 15 fractions (0.74 ml/ fraction) were collected on a BIOCOMP gradient fractionator.

For RT-qPCR, total RNA was isolated from 250 μl of each gradient fraction using TRIzol LS reagent. cDNA was generated from equal amount of RNA by reverse transcription using the TransScript All-in-One First-Strand cDNA Synthesis SuperMix for qPCR. The relative quantity of specific mRNAs was measured by quantitative polymerase chain reaction (qPCR) using the TransStart Top Green qPCR SuperMix with the Real-Time PCR System. Primers for qPCR are listed in the **Table V1**. Total ribosome occupancy was calculated according to the method of (Darnell et al., 2011). Percentage of mRNA in each fraction as determined by qPCR was multiplied by the number of ribosomes in that fraction (extrapolated from UV traces based on the linearity of the sucrose density gradient) and summed over the gradient.

### Pulsed SILAC

Cells were grown in McCoy’s 5A media (SILAC standard, 88441) containing light (^12^C, ^14^N) lysine and arginine supplemented with 10% dialyzed FBS for 2 weeks. 2 × 10^6^ cells were treated with 500 µM IAA or DMSO for 6 h before switching to heavy medium containing (^13^C, ^15^N) lysine and arginine supplemented with 500 µM IAA. After another 6 h, cells were scraped in pre-cooled PBS and pelleted. Cell pellets were resuspended in 0.5 ml 8 M urea lysis buffer (8 M urea in 100 mM Tris/HCl, pH 8.0). Following centrifugation at 12,000 g for 15 min at 4°C, supernatants were collected, and protein concentrations were determined by BCA assay. 200 ug protein was loaded onto a Microcon Centrifugal Filters (Millipore, Cat: 42407). Filter aided sample preparation (FASP) for LC-MS/MS was done as described (Wiśniewski et al., 2011). The digested peptide mixtures were re-dissolved in 0.1% formic acid in ultrapure water, and LC-MS/MS was performed as described (Lin et al., 2020). Protein identification and quantitation were performed with Thermo Proteome Discoverer (PD 2.1.1.2.) software searching against the UniProt human protein database release 2018_04. Triplicate LC-MS/MS datasets were subjected to statistical testing (Benjamini-Hochberg) to identify proteins whose synthesis (Log2 Heavy IAA/Heavy DMSO) was changed upon depletion of eIF3 subunits at an FDR ≤ 0.1.

### Quantitative proteomics of eIF3 complexes

eIF3-mAID cell lines were grown in McCoy’s 5A media (SILAC standard, 88441) containing light (^12^C, ^14^N) lysine and arginine or containing heavy (^13^C) lysine and arginine for 2 weeks. 12 h prior to harvest, 500 µM IAA or vehicle (DMSO) were added. Cells were collected and lysed in IP buffer (20 mM Tris/HCl pH 7.5, 150 mM NaCl, 0.5% Triton X-100 and 1 x Pierce™ protease inhibitor cocktail) through rotation for 15 min at 4°C. Supernatants were collected after centrifugation at 13,000 g for 15 min at 4°C, and protein concentrations were determined by BCA assay. Lysates were split into two equal parts and 2 μg antibodies (anti-eIF3b or anti-eIF3c) were added and incubated by rotating overnight at 4°C. 50 μl Protein A Surebead, (BIO-RAD, #1614813) were added to the mixture. Following incubation at 4°C for 4h with constant rotation, beads were washed with 500 μl IP buffer 3 times 10 min each. Following trypsin digestion, corresponding heavy and light samples were mixed at a ratio of 1:1. After desalting, peptide mixtures were re-dissolved in 0.1% formic acid in ultrapure water, and LC-MS/MS, protein identification and quantitation were performed as described above.

### RNA extraction and RNA sequencing

Total RNA and RNA contained in heavy polysomal fractions were isolated using the Trizol reagent (Life Technology). Library preparation for bulk-sequencing of poly(A)-RNA was done as described previously (Parekh et al., 2016). Briefly, barcoded cDNA of each sample was generated with a Maxima RT polymerase (Thermo Fisher) using oligo-dT primer containing barcodes, unique molecular identifiers (UMIs) and an adaptor. 5’-Ends of the cDNAs were extended by a template switch oligo (TSO) and full-length cDNA was amplified with primers binding to the TSO-site and the adaptor. NEB UltraII FS kit was used to fragment cDNA. After end repair and A-tailing a TruSeq adapter was ligated, and 3’-end-fragments were finally amplified using primers with Illumina P5 and P7 overhangs. In comparison to (Parekh et al., 2016), the P5 and P7 sites were exchanged to allow sequencing of the cDNA in read1 and barcodes and UMIs in read2 to achieve a better cluster recognition. The library was sequenced on a NextSeq 500 (Illumina) with 57 cycles for the cDNA in read1 and 16 cycles for the barcodes and UMIs in read2. Data was processed using the published Drop-seq pipeline (v1.0) to generate sample- and gene-wise UMI tables (Macosko et al., 2015). Reference genome (GRCh38) was used for alignment. Transcript and gene definitions were used according to the GENCODE version 38.

### RNA seq data analysis

Only genes with at least one raw read in all samples were retained in the analysis. Raw RNA-seq datasets were normalized for sequencing depth by calculating counts per million mapped reads (CPM) using the EdgeR package (version 3.38.0) in R. For polysomal RNA samples, an additional normalization step was performed according to the formula CPM x (P_IAA_ / P_DMSO_) with P_x_ signifying the area under the curve of the polysomal fraction of the corresponding lysates separated by sucrose gradient density centrifugation. This normalization was done to account for the different levels in total polysomes in the DMSO and IAA treated sample. This was particularly important for those cell lines that showed a strong decrease in polysomes, including eIF3a-mAID, eIF3e-mAID, and eIF3f-mAID; but for consistency, the normalization was performed for all 6 cell lines. In the subsequent step, edgeR (Robinson et al., 2010) was used to identify changed mRNAs. mRNAs were considered changed in either total or polysome RNAseq if they differed between the DMSO-treated and IAA-treated samples (Log2 IAA/DMSO) with FDR ≤ 0.05. Translational efficiencies (TEs) were calculated as Log2 [(polysomal mRNA IAA / total mRNA IAA) / (polysomal mRNA DMSO / total mRNA DMSO)] for data with cpm ≥ 10. Significance was determined with the R-package’s p.adjust function with the FDR≤ 0.2.

### RNA immunoprecipitation and qPCR

70-80% density cells were cultured in McCoy’s 5A in 150 mm dishes and lysed in 800 µl RNA immunoprecipitation (RIP) lysis buffer containing 20 mM Tris-HCl pH7.5, 130 mM KCl,10 mM MgCl_2_,1 mM EDTA, 0.5% NP-40, 0.5% sodium deoxycholate, 0.5 mM DTT supplemented with 1 x Pierce protease inhibitor cocktail, 100 U/ml RNase inhibitor, 0.4 U/ml DNase. The lysate was incubated on ice for 15 min. The supernatants were collected after centrifugation at 13,000 g for 15 min at 4°C, and protein concentrations were determined by BCA assay. 3 ug rabbit IgG or anti-eIF3c antibody were attached to 20 µl Protein A Surebead, (BIO-RAD, #1614813) in RIP lysis buffer for 30 min at room temperature. Beads were washed three times in RIP lysis buffer and added to 1.2 mg protein lysate followed by rotation for 4 h at 4 °C. Beads were washed four times for 5 min at 4 °C with RIP washing buffer containing 0.1% SDS, 1% Triton-X 100, 2 mM EDTA, 20 mM Tris-HCl pH 8.0, 500 mM NaCl. Bound RNA was isolated from the washed beads by adding Trizol and quantified by qPCR as described above.

### Functional pathway analysis

For each given gene list, pathway and process enrichment analysis has been carried out with Metascape (metascape.org (Zhou et al., 2019)) with the following ontology sources: GO Biological Processes, GO Cellular Components. The 9895 human proteins detected by LC-MS/MS in all combined datasets were used as the enrichment background. Terms with a p-value < 0.01, a minimum count of 3, and an enrichment factor > 1.5 (the enrichment factor is the ratio between the observed counts and the counts expected by chance) were collected and grouped into clusters based on their membership similarities. According to the description at metascape.org, p-values were calculated based on the accumulative hypergeometric distribution, and q-values were calculated using the Benjamini-Hochberg procedure to account for multiple testing.

### Computational modeling

We created a dynamical model, which consists of a network of a ribosome flow model (RFM) and a pool of free ribosomes (i.e. ribosomes that are not attached to any mRNA molecule) to analyze the effect of the different eIF3 subunits on translation similar to the approach used in (Zarai and Tuller, 2018). The RFM is a deterministic mathematical model that can be used to model and analyze many transport phenomena (Reuveni et al., 2011). It is based on a simple exclusion principle of n compartments (or sites, e.g., a site along the mRNA, gene, microtubule) with n state variables describing the compartments particle density and n dynamical equations describing the flow dynamics between these compartments as a function of time. The flow between consecutive compartments is controlled by λ_i parameters. Specifically, λ_0 denotes the initiation rate, and λ_i,i∈{1,…n}, denotes the elongation rate from site i to site i+1. These rates depend on various factors including availability of tRNA molecules, amino acids, aminoacyl tRNA synthetase activity and concentration, and local mRNA folding. We denote by R the steady state output flow, and by e_i,i=1,…n, the steady-state particle density at site i. In the context of mRNA translation, R is the steady state translation rate (i.e., the steady state protein production rate), and e_i is the steady state ribosomal density at site i.

As a specific example, we modeled the translation of the gene TP53 (transcript ENST00000269305.9), which contains 393 amino acids. We divided the mRNA to non-overlapping pieces. The first piece includes the first 9 codons that are related to various stages of initiation (Tuller and Zur, 2015). The other pieces include 10 non-overlapping codons each, except for the last one that includes between 5 and 15 codons. We model every piece of the mRNA as a RFM site, resulting in 38 sites. We estimated the elongation rates λ_i at each site using ribo-seq data from HEK293 cells for the codon decoding rates (Dana and Tuller, 2015), normalized so that the median elongation rate becomes five codons per second. We note that similar modeling can be performed for any transcript. Let *z* denote the free ribosomal pool, *v*_*k*_ the eIF3k concentration value, and let

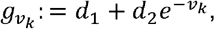

where *d*_1_ >0 and *d*_2_ > 0 are constant parameters. We model *z* as

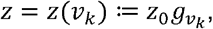

where *z*_0_ denotes the free ribosomal pool governed by the dynamics with the RFM (see below), and 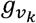 models the effect of *v*_*k*_ on the free ribosomal pool.

Let *f* (*v*) = *c*_1_ + *c*_2_ (1 − *e*^*-v*^), for all *v* ≥ 0, and for any positive parameter set *c*_1_, *c*_2_. We use *f* (*v*) to model the effect of each eIF3 a, b, e, and f subunit on the translation initiation rate, where v denotes the corresponding subunit concentration value. The parameter *c*_1_ and *c*_2_ can generally vary between the a, b, e, and f subunit models, however, for simplicity they are assumed to have the same values for all these models in the analysis.

Let

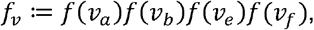

denote the product of the subunits a, b, e, and f models.

Let *x*_*i*_ = *x*_*i*_ (*t*) denote the ribosomal density at site (in the RFM) at time *t*, 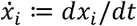, and

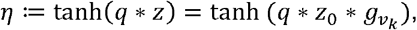

where *q* > 0 is a normalization factor, and tanh () is the hyperbolic tangent function. The usage of this function is derived from (Zarai and Tuller, 2018).

The equations of our dynamical model, which consists of a network of a RFM and the free ribosomal pool and is based on (Zarai and Tuller, 2018), are given by

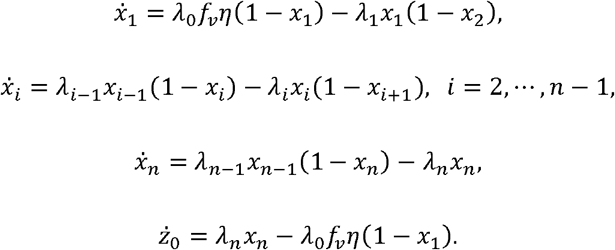

This dynamical model admits a unique steady state for a given total number of ribosomes in the network (Raveh et al., 2016). The concentrations of the eIF3 a, b, e, and f subunits affect the effective initiation rate by the product *λ*_0_*f*_*v*_. eIF3k concentration affects the free ribosomal pool, which in turn affects the effective initiation rate by the product ; *λ*_0_*f*_*v*_*η*. The dynamical model parameter values used in the simulations were adjusted so that 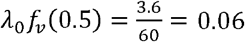 (Yan et al., 2016) and are given by: *c*_1_ = 0.59, *c*_2_ = 1.11, *d*_1_ = 0.65, *d*_2_ = 600.0, and *q* = 1.2 *10^−5^. Given these parameter values, the steady state free ribosomal pool at 0.5 concentration of each eIF3 subunit is 28,293.

### Quantification and statistical analysis

All FDR and p values for the RNAseq data were calculated using EdgeR in R. Statistical analysis of the remaining datasets (proteomic and TE) was performed with Microsoft Excel. P values for the remaining datasets were determined using the multiple t-tests function (two-stage step-up method of Benjamini, Krieger, and Yekutieli) and unpaired Student’s t-test in GraphPad Prism 8.

## Supporting information

Reagents_Tools_Table

## Data availability

RNA-seq data were deposited into European Nucleotide Archive (accession number: PRJEB53197). Proteomic data were deposited to ProteomeXchange via PRIDE (accession number: PXD034084).

## ACKNOWLEDGEMENTS

We thank R. Ding and the Mass Spectrometry Facility of the School of Pharmaceutical Sciences at Xiamen University for support. The support from the Equipment Platform of the State Key Lab of Cellular Stress Biology at Xiamen University is gratefully acknowledged. This work was funded through grant grants 81773771 and 31770813 from the National Science Foundation of China (D.A.W), the Natural Science Foundation of Fujian Province (2018J01053), and the Innovation Program of Xiamen University Department of Life Sciences & Human Health (Y.C.). R.R. is supported by the European Research Council (Consolidator Grant PACA-MET and MSCA-ITN-ETN PRECODE), the Deutsche Forschungsgemeinschaft (DFG RA1629/2-1; DFG RA1629/4-1; SFB1321; SFB1335), the German Cancer Consortium, and the Deutsche Krebshilfe (70114314). Additional funding was provided by the Zimin Institute for Engineering Solutions Advancing Better Lives (T.T.).

## AUTHOR CONTRIBUTIONS

Conceptualization, H.D. and D.A.W.; Methodology, H.D., S.Z., Y.W., L.S., G.T., C.H., Y.Z., and R.Ö.; Formal Analysis, H.D., S.Z., Y.W., L.S., Y.Z.; Investigation, H.D., S.Z., Y.W., L.S., G.T., C.H., Y.Z., and R. Ö.; Writing – Original Draft, H.D. and D.A.W.; Writing – Review & Editing, H.D., S.Z., Y.Z., Y.W., L.S., G.T., C.H., R.Ö., R.R., Y.C., T.T., and D.A.W.; Visualization, H.D., Y.Z., D.A.W.; Supervision, R.R., T.T., Y.C. and D.A.W.; Funding Acquisition, R.R., T.T., Y.C. and D.A.W.

## COMPETING INTERESTS

The authors declare no competing financial interests.

## FIGURE LEGENDS

**Supplementary Figure EV1.**
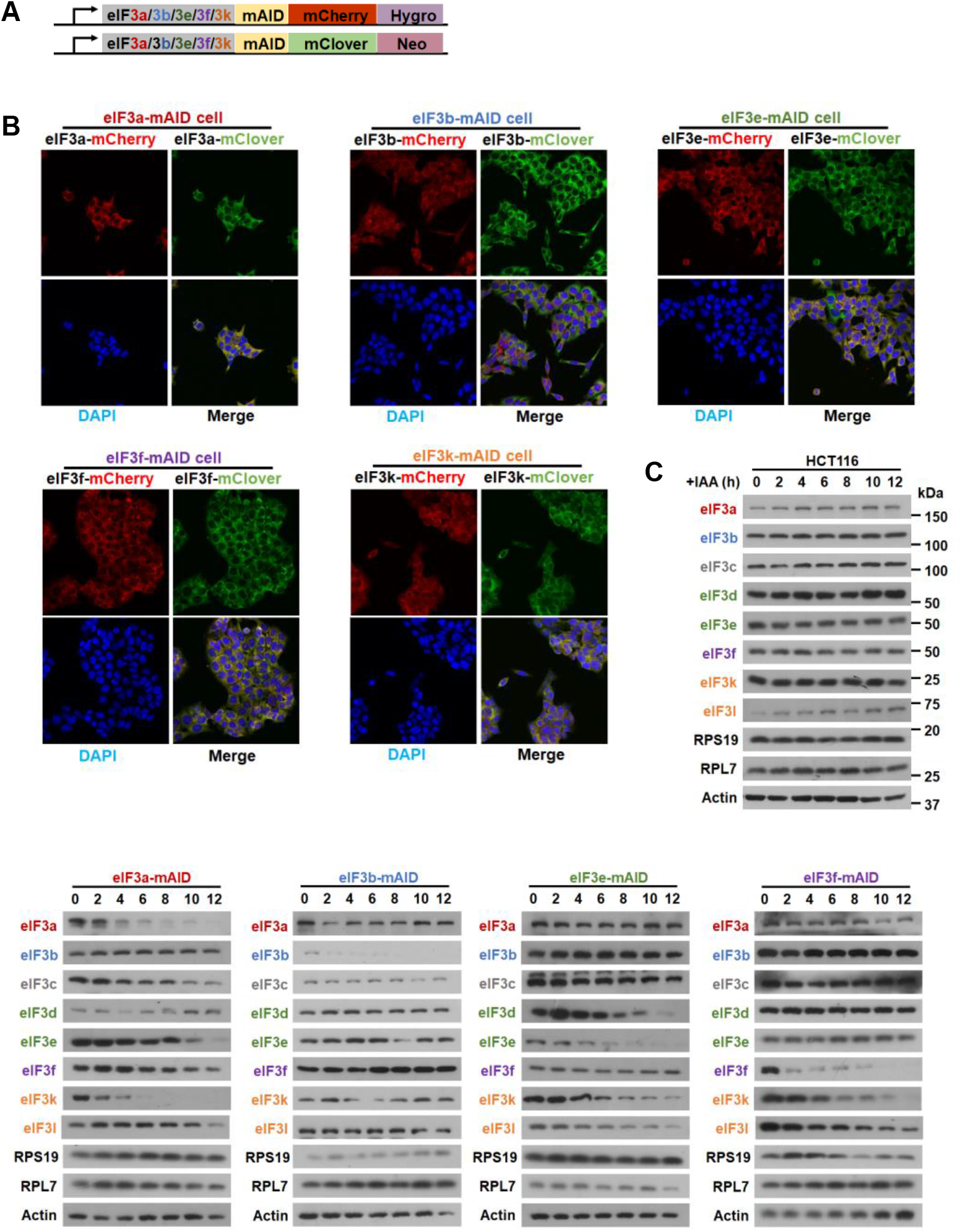
Characterization of HCT116-derived cell lines engineered for inducible degradation of eIF3 subunits (refers to Figure 1) **(A)** Schematic of the conditional alleles of eIF3 subunits. Both alleles of selected eIF3 genes were tagged with the auxin-inducible degradation domain (mAID). In addition, single alleles were tagged either with mClover using neomycin as the selection marker or with mCherry using hygromycin (Natsume et al., 2016). **(B)** Life cell fluorescence micrographs of the indicated eIF3-mAID cell lines. **(C)** The indicated eIF3-mAID cells lines were exposed to 500 μM indole-3-acetic acid (IAA) for the periods shown, and cell lysates were subjected to immunoblotting with the indicated antibodies.

**Supplementary Figure EV2.**
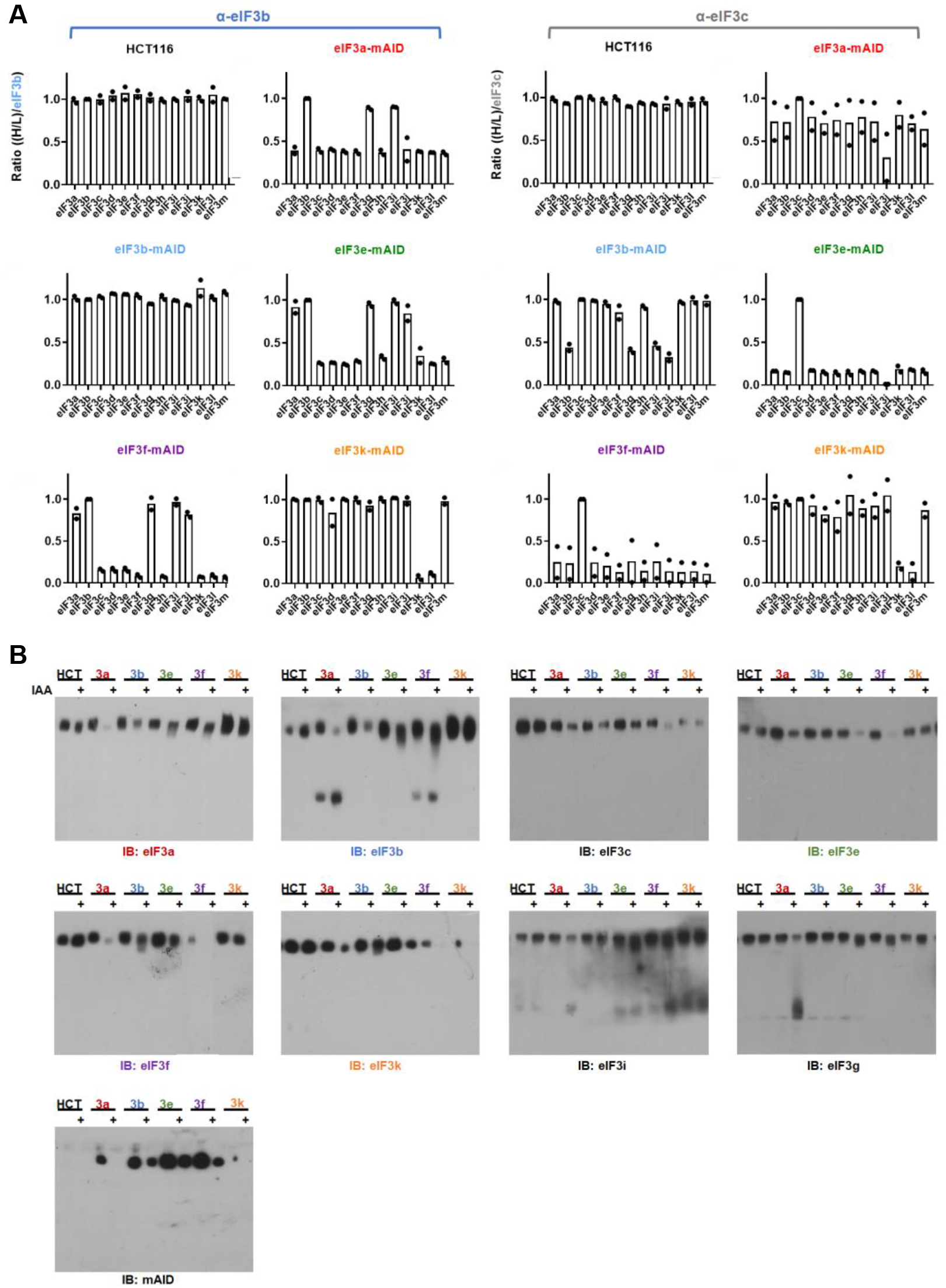
Effect of conditional depletion of eIF3 subunits on the eIF3 holo-complex (refers to Figure 1) **(A)** The graphs show the duplicate data condensed in the heatmap shown in Figure 1C. **(B)** Native PAGE analysis of eIF3 in the indicated cell lines. Cells were exposed to IAA for 12 h, cell lysates were separated by native PAGE, and the abundance and integrity of eIF3 was probed with antibodies directed against the indicated subunits.

**Supplementary Figure EV3.**
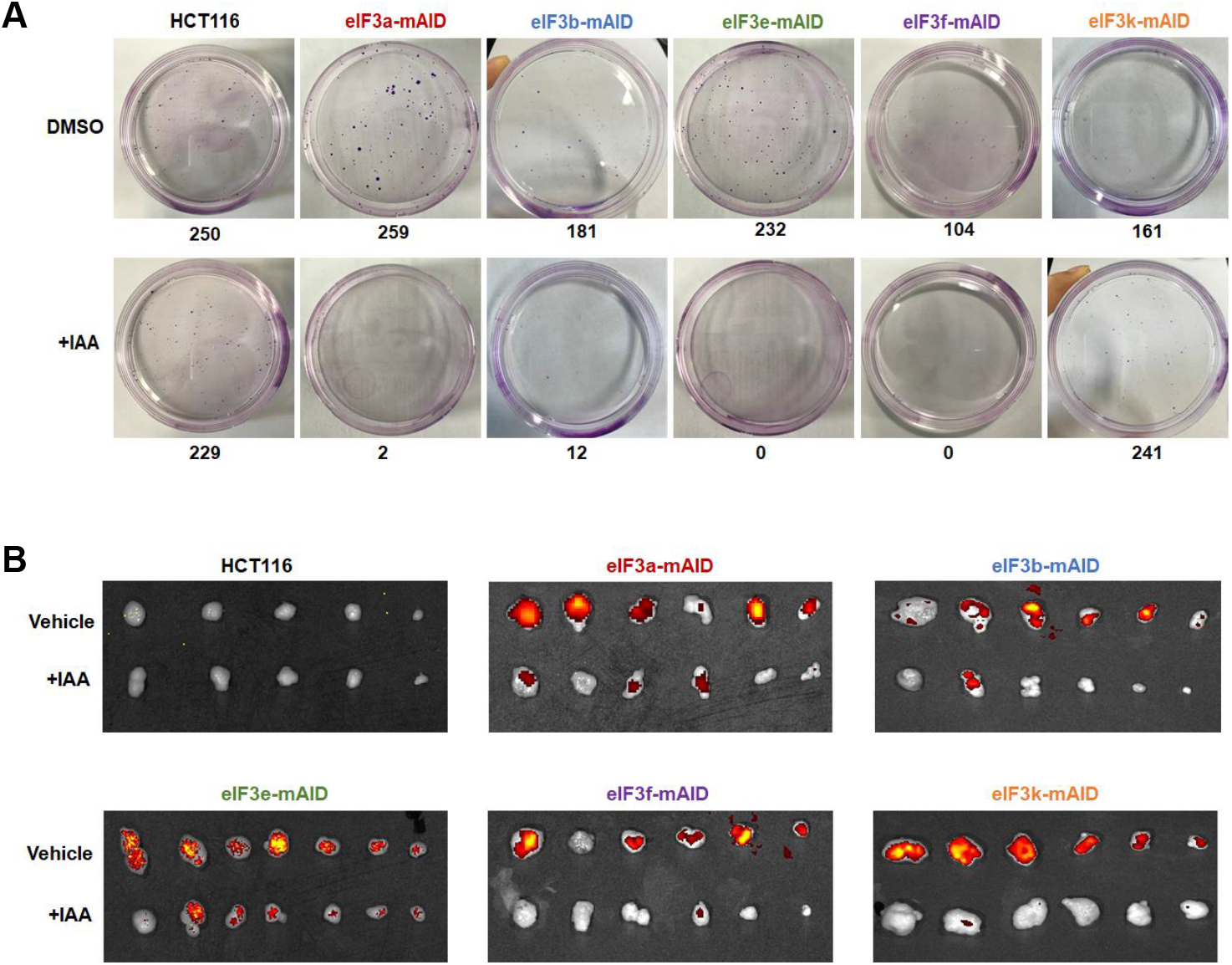
Effect of depletion of eIF3 subunits on cell and tumor growth (refers to Figure 2) **(A)** Representative plates for the colony formation experiment shown in Figure 1C. 1000 cells of the indicated cell lines were plated and grown in DMSO or IAA for 2 weeks. Colonies were fixed, stained, and counted. **(B)** Fluorescence imaging of the tumors grown from the indicated cell lines. The images document strong downregulation of mClover-tagged eIF3 subunits in tumors from mice treated with IAA.

**Supplementary Figure EV4.**
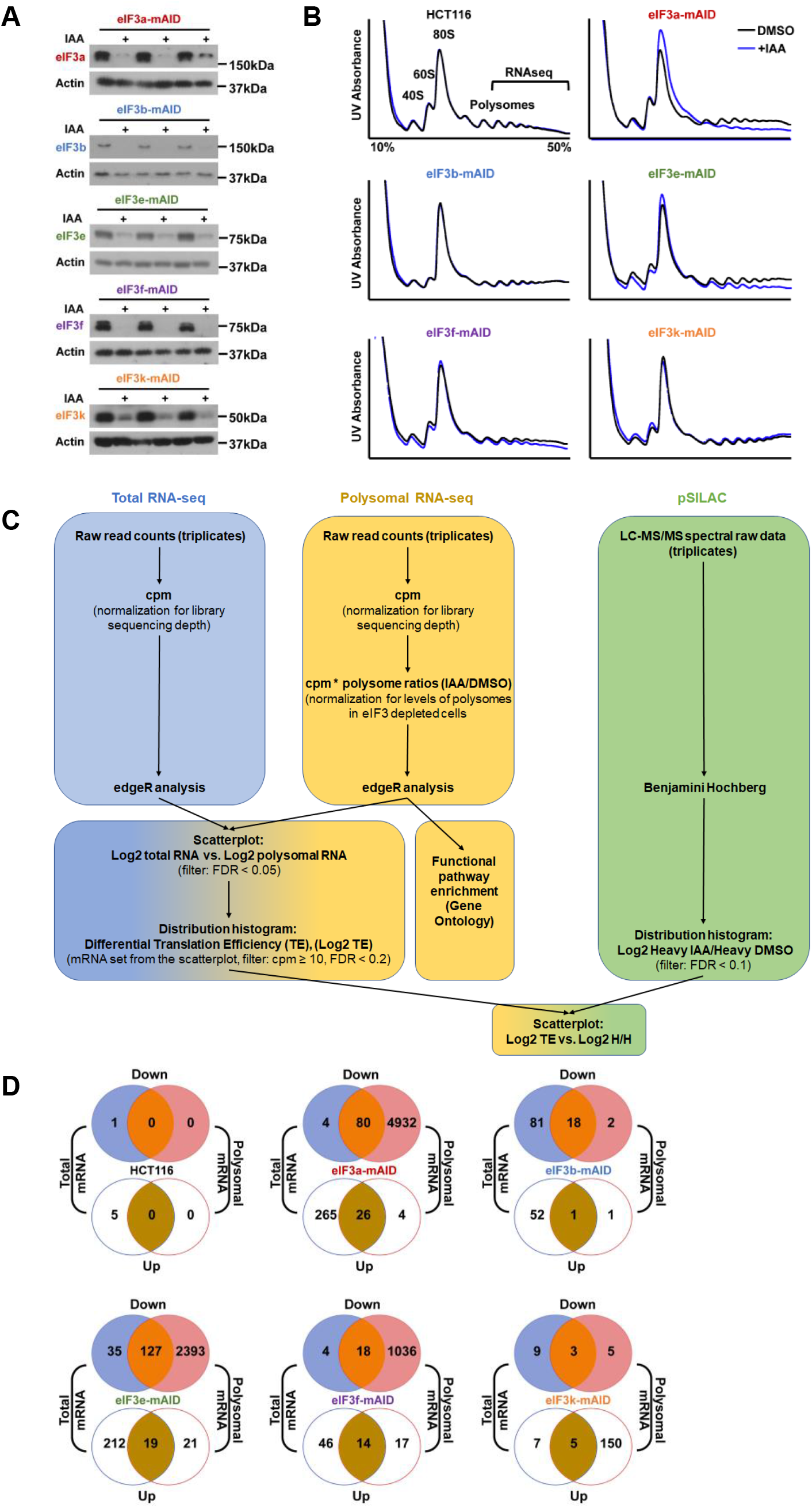
Effect of depletion of eIF3 subunits on mRNA translation (refers to Figure 3) **(A)** Primary data for the graph shown in Figure 3A. The indicated conditional eIF3 cell lines were exposed to 500 μM IAA for 12 h to induce the degradation of the respective eIF3 subunit. Cell lysates were subjected to immunoblotting to quantify the expression of the indicated eIF3 subunits. Actin signal was used as a reference. **(B)** Representative primary data for the graph in Figure 3B. The indicated cell lines were exposed to IAA for 12 h, and cell lysates were separated by sucrose density gradient centrifugation. The monosomal (40S, 60S, 80S) and polysomal peaks were quantified, and P/M ratios relative to the vehicle DMSO are plotted in Figure 3B. The region of the gradient used for extraction of polysomal RNA for RNA-seq is indicated. **(C)** Strategy of analyzing transcriptomic and proteomic datasets. Total (blue panel) and polysomal (yellow) RNA samples were collected from triplicate cultures of eIF3-mAID cells exposed to DMSO or IAA for 12 h. Raw RNA-seq datasets were normalized for sequencing depth by calculating counts per million mapped reads (CPM). For polysomal RNA samples, an additional normalization step was performed to account for the different levels in total polysomes in each sample. This was particularly necessary for those cell lines that showed a strong decrease in polysomes, including eIF3a-mAID, eIF3e-mAID, and eIF3f-mAID; but the normalization was performed for all 6 cell lines. In the next step, edgeR (Robinson et al., 2010) was used to identify changed mRNAs. mRNAs changed in polysomes (FDR ≤ 0.05) were analyzed for enriched Gene Ontology functional terms (Figure 3D). mRNAs changed (FDR ≤ 0.05) in the total and polysomal populations were then graphed in a scatter plot (Figure 3C). Translational efficiencies (TEs) were calculated for all data points included in the scatter plot with cpm ≥ 10 as Log2 [(polysomal mRNA IAA / total mRNA IAA) / (polysomal mRNA DMSO / total mRNA DMSO)], and TEs were plotted in histograms (Figure 3E). Triplicate LC-MS/MS datasets were subjected to statistical testing (Benjamini-Hochberg) to identify proteins whose synthesis (Log2 Heavy IAA / Heavy DMSO) was changed upon depletion of eIF3 subunits at a FDR ≤ 0.1. Log2 Heavy IAA / Heavy DMSO values were graphed in a histogram (Figure 3F). Changes in TE and protein synthesis were compared in a scatter plot of Log2 TE versus Log2 Heavy IAA/Heavy DMSO (Figure S5E). **(D)** Venn diagrams summarizing the changes in the total and polysomal mRNAs (FDR ≤ 0.05) in the indicated cell lines.

**Supplementary Figure EV5.**
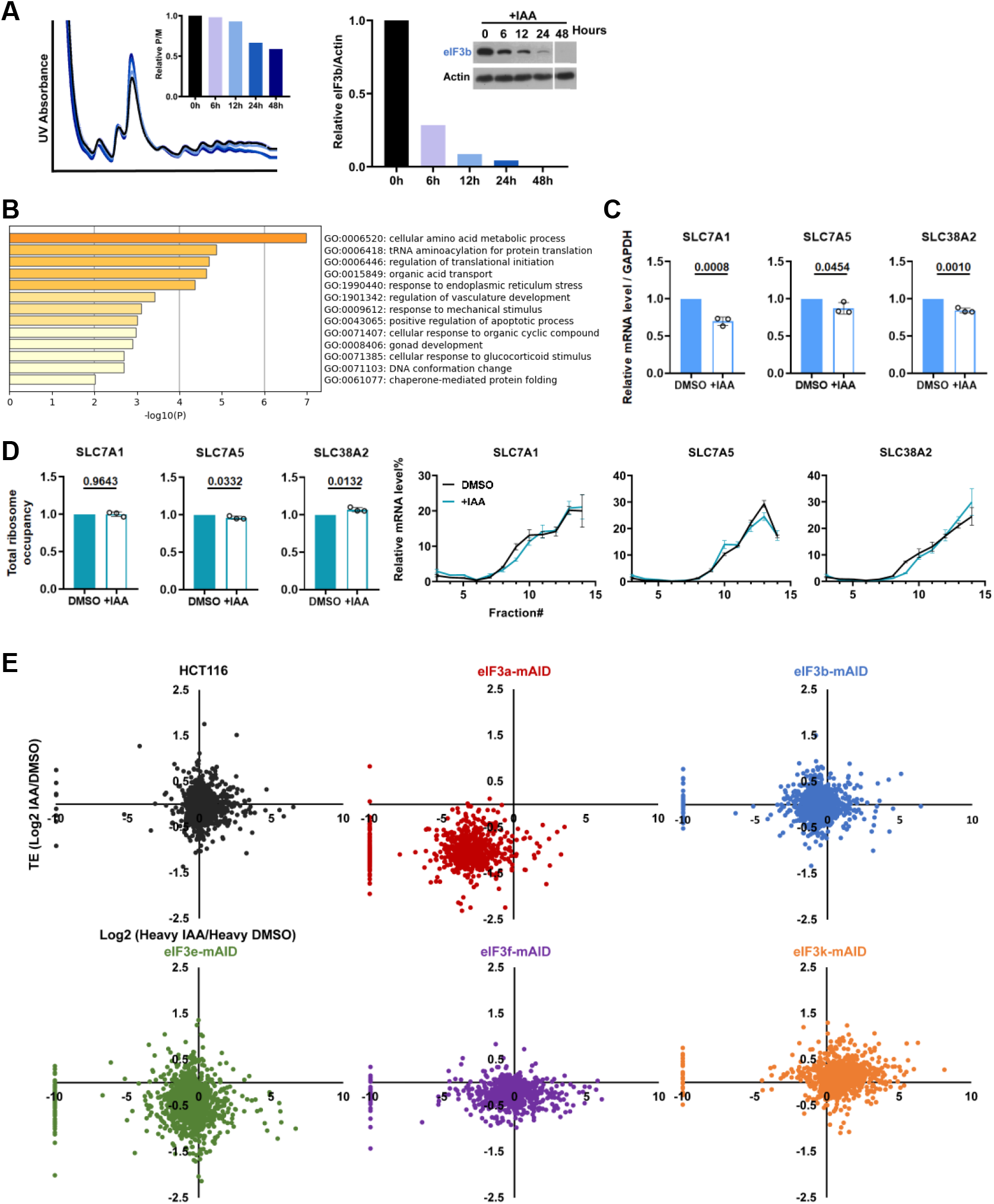
Effect of depletion of eIF3b on translation and global correlations between TE and protein synthesis (refers to Figure 3) **(A)** eIF3b-mAID cells were exposed to IAA for the indicated periods and cell lysates were separated by sucrose density gradient centrifugation. Polysome/monosome ratios were determined and displayed in a bar graph (left panel). The expression of eIF3b in the lysates was quantified by immunoblotting, and results are displayed in a bar graph (right panel). **(B)** The set of 81 mRNAs downregulated in the total but not the polysomal mRNA fraction upon depletion of eIF3b was examined for enriched GO terms. **(C)** RT-qPCR analysis of the levels of the indicated mRNAs in total RNA prepared from eIF3b-mAID cells exposed to DMSO or IAA for 12 hours. Data were normalized to the signal obtained for GAPDH. Bars represent means ± SD, n = 3; numbers indicate p values (unpaired Student’s t-test). **(D)** Total ribosome occupancy of the indicated mRNAs in eIF3b-mAID cells exposed to DMSO or IAA for 12 hours was determined by RT-qPCR of RNA across a sucrose density gradient (see Materials and Methods). Bars represent means ± SD, n = 3; numbers indicate p values (unpaired Student’s t-test). **(E)** Scatter plots of changes in TE versus protein synthesis (Log2 Heavy IAA/Heavy DMSO) upon depletion of the indicated eIF3 subunits.

**Supplementary Figure EV6.**
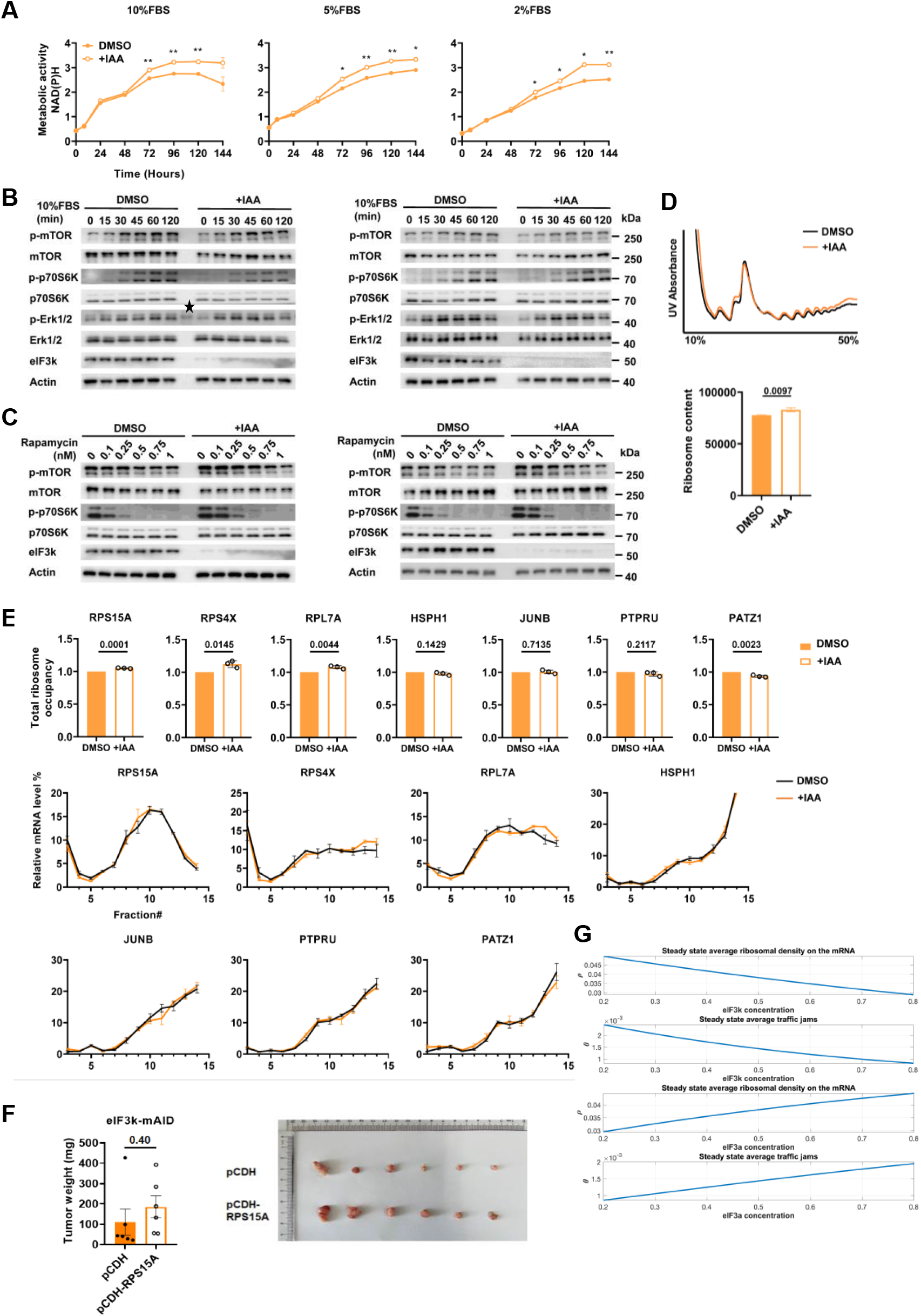
Effect of eIF3k depletion on growth factor sensitivity, ribosome content, and RPS15A (refers to Figures 4 and 5) **(A)** eIF3k-mAID cells were maintained in media containing different concentrations of FBS and DMSO or IAA as indicated, and metabolic activity (i.e. NAD(P)H) as a proxy of cell proliferation was determined by CCK-8 assay. Data represent means ± SD, n = 3. Asterisks denote: * p < 0.005, ** p < 0.00005, *** p < 0.000005 (two-stage step-up method of Benjamini, Krieger, and Yekutieli). **(B)** eIF3k-mAID cells were serum starved by maintaining in media containing 0% FBS for 24 h. During the last 12 h of starvation, DMSO or IAA was added as indicated. Cells were re-stimulated with media containing 10% FBS, and the expression of the indicated proteins was followed of a period of 120 minutes. Duplicate experiments are shown (quantification in Figure 4B). The asterisk denotes a size marker band. **(C)** eIF3k-mAID cells were exposed to IAA or DMSO for 12 hours, followed by the addition of increasing concentrations of rapamycin for 1 h. The expression of the indicated proteins was determined by immunoblotting. Duplicate experiments are shown (quantification in Figure 4C). **(D)** Lysates of eIF3k-mAID cells exposed to DMSO or IAA for 48 h were separated by sucrose density gradient centrifugation. Total ribosome content was determined by integrating the monosomal and polysomal peaks and plotted. Error bars represent means ± SD, n = 3. Numbers indicate p values (unpaired Student’s t-test). **(E)** Total ribosome occupancy of the indicated mRNAs in eIF3k-mAID cells exposed to DMSO or IAA for 48 hours was determined by RT-qPCR of RNA across a sucrose density gradient (see Materials and Methods). Bars represent means ± SD, n = 3; numbers indicate p values (unpaired Student’s t-test). Triplicate RT-qPCR data across the sucrose gradient are shown below the bar graphs. **(F)** 1 × 10^6^ eIF3k-mAID cells stably expressing ectopic RPS15A (pCDH-RPS15A) or empty vector (pCDH) were injected into nude mice and tumor growth was followed for 2 weeks. Graphs represent means of final tumor weights ± SD, n = 6. Numbers indicate p values (unpaired Student’s t-test). **(G)** Simulated effect of eIF3k and eIF3a concentration on the average steady-state density and traffic jam. The concentration of other eIF3 subunits is kept constant at 0.5.

**Supplementary Figure EV7.**
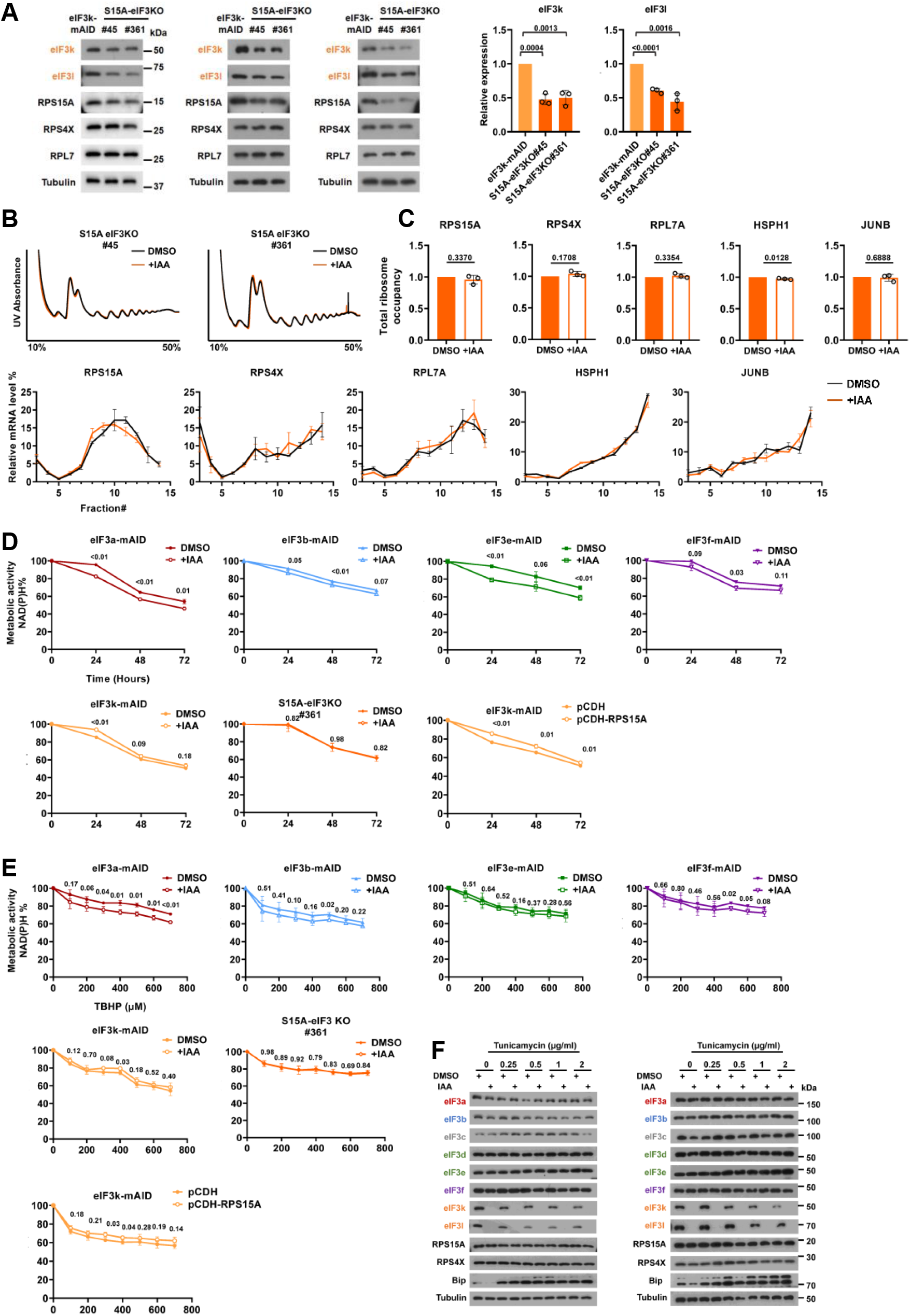
Effect of the eIF3 binding site on cell proliferation, tumor growth, RPS15A, and stress sensitivity (refers to Figures 6 and 7) **(A)** Basal expression of the indicated proteins was determined in parental eIF3k-mAID cells or S15A-eIF3KO cells (clones #45 and # 361) by immunoblotting. Triplicate experiments are shown (quantification in Figure 6E). **(B)** Lysate of S15A-eIF3KO cells (clones #45 and # 361) exposed to DMSO or IAA for 12 h were separated by sucrose density gradient centrifugation. Representative example of triplicate data used for quantification of total ribosome content in Figure 6H. **(C)** Total ribosome occupancy of the indicated mRNAs in S15A-eIF3KO (clone #45) cells exposed to DMSO or IAA for 12 hours was determined by RT-qPCR of RNA across a sucrose density gradient (see Materials and Methods). Bars represent means ± SD, n = 3; numbers indicate p values (unpaired Student’s t-test). Triplicate RT-qPCR data across the sucrose gradient are shown below the bar graphs. **(D)** The indicated cell lines were treated with DMSO or IAA for 12 hours, followed by exposure to 2 µg/ml tunicamycin for up to 72 hours. Metabolic activity (i.e. NAD(P)H) as a proxy of cell viability was determined by CCK-8 assay at the times indicated. Graphs represent means ± SD, n = 3. Numbers indicate p values (two-stage step-up method of Benjamini, Krieger, and Yekutieli). **(E)** Same experiment as in (D) but cells were exposed to increasing concentrations of the oxidative stress inducer tert-butyl hydroperoxide (TBHP) for 2 hours. **(F)** The indicated eIF3-mAID cell lines were treated with DMSO or IAA for 12 hours, followed by exposure to 2 µg/ml tunicamycin for 24 hours. The expression of individual eIF3 subunits, ribosomal proteins, and the ER stress marker BIP were assessed by immunoblotting. Tubulin is show for reference. Replicates for the data quantified in Figure 7C.

## Notes

### Competing Interest Statement

The authors have declared no competing interest.

### Summary of Updates

The author order revised.

